# A new isolated local varicella virus: isolation, identification, comparative growth characteristics and immunological evaluation on an animal model

**DOI:** 10.1101/2020.12.03.409672

**Authors:** Fatemeh Esna-Ashari, Abbas Shafyi, Farzaneh Sabahi, Mehrdad Ravanshad, Zohreh-Azita Sadigh, Ashraf Mohammadi

**Author notes:** **Corresponding Author: Ashraf Mohammadi:** Associate Professor Scientific member, Head of Human Viral Vaccine Department, Razi Vaccine & Serum Research Institute, Agricultural Research, Education & Extension Organization (AREEO), Karaj, I.R.Iran. Int.2336. **Data Availability:** All relevant data are within the paper and its Supporting Information files. **Competing interests:** The authors have declared that no competing interests exist. **Other author Fatemeh Esna-Ashari:** Ph.D. candidate of medical virology, Department of Virology, Faculty of Medical Sciences, Tarbiat Modares University, Tehran, I.R. Iran.;, **Professor Abbas Shafyi:** Scientific member of Human Viral Vaccine Department, Razi Vaccine & Serum Research Institute, Karaj, I.R.Iran., **Professor Farzaneh Sabahi:** Ph.D., Scientific member, Department of virology, faculty of medical sciences, Tarbiat Modares University, Tehran, I.R. IRAN. **Dr. Mehrdad Ravanshad:** Ph.D., Scientific member, Department of virology, faculty of medical sciences, Tarbiat Modares University, Tehran, I.R. IRAN. **Dr.Zohreh-Azita Sadigh:** Ph.D., Scientific member of Human Viral Vaccine Department, Razi Vaccine & Serum Research Institute, Karaj, I.R.Iran.

## Abstract

A panel of 4 different cell lines was optimized for isolation, identification, and authentication of a VZV virus from a swab sample of an 8-year-old boy suspected to varicella zoster infection. The system enabled highly efficient and rapid isolation of viruses in 33°C by serial sub culturing to more than 25 passages. The technique relies on isolation of viral genes by increasing the number of particles that are statistically represented in cell culture and verified by CCID_50_, FAM-RT-PCR, and IE62 antibody in IF test. The viral genes (ORF38, ORF54) confirmed the new isolate as VZV and revealed the amino acid sequence of viral-encoded proteins after 27 passages, identical with positive control virus, in RFLP-PCR test. Utilization of successive serial passages at temperatures lower than the normal body temperature would reduce the virus virulence and directly cause virus attenuation. As a result, the attenuated virus is adapted to growth in vitro and presented higher replication at 33°C. Our goal was to determine if the targeted virus with a large double-stranded DNA genome, varicella virus, is isolated and can be attenuated by cold adapting in vitro, Using vOka as attenuated VZV golden standard, in two quantitative tests, including CCID50 and FAM-RT-PCR. Finally, when compared with the local isolated virus, these results were strongly confirmed. We recorded plaque forming assay to show phenotypic changing which encodes the attenuation regarding size of plaque. Although in plaque forming assay, the size of plaque seemed smaller at first glance, the statistical distribution of the plaque size did not show any change between the virus in the first and last passages. In cell culture, the local VZV isolated viruses formed clear plaques and grew to higher titers compared with lower passages as parental virus.Due to lack of access to human fetal lung cells (MRC-5) and an alternative to vaccine production in the future, a new authenticated local foreskin cell substrate (RFSC) was used for virus cultivation. In comparison to, MRC-5-optimized and cloned-viruses replicated in vitro with kinetics that were similar to those of the RFSC. Laboratory animals that were infected with the optimized virus fluid as vaccine showed a good neutralization antibody against local VZV isolated as compared to vOka as control positive virus. These results demonstrate that the virus isolated from swab sample was authenticated as VZV virus, and this cold adapted attenuated virus may be an applicable candidate for future plan.

**Author summary:** We used different cell substrates for isolation, identification, and attenuation of a new local VZV virus from vesicle swab of suspected patient to varicella zoster diseases. The technique involves serial sub culturing of virus in cell culture. Our goal was to determine if the targeted virus with a large double-stranded DNA genome, varicella virus, is isolated and can be attenuated by cold adapting in vitro, using vOka as attenuated VZV golden standard, The virus was cloned and purified by serial dilution cloning and plaque assay. Different techniques were used regarding verification as differentiation from other members in family, potency, and sterility for isolated virus. The virus can grow well in new local foreskin cell substrate and produce good titre as compared with MRC-5. The isolated VZV potentially induced neutralization antibody in animal model. The results of our study imply that local VZV might be an applicable isolate for future plan in research and developing a varicella vaccine.

## Introduction

Infectious pathogens are a serious problem in human life and varicella Zoster Virus (VZV) is a particularly challenging germ behind the disease. Varicella “Human Herpes Virus type 3 (HHV3)” is a cell-associated DNA virus belong to Alpha-herpes viridae. The virus causes varicella, a lymph proliferative disease with severe clinical symptoms in human. This is an acute infection which is characterized by fever, malaise, headache, and abdominal pain 1-2 days before appearance of Macula popular rash. During several days’ rash involves 3 or more successive crops; each crop usually progresses within less than 24 h from macules to papules, vesicles, pustules, and crusts. In this case, there are lesions in different stages of development on any part of the body [1]. These clinical signs usually start on face and trunk, then spread to extremities. These features involve 250-500 lesions that are pruritic. The lesions are typically crusted 4-7 days after rash onset, and affects both children and young adults. Varicella disease is classified based on the virulence exhibited in infection through unvaccinated or vaccinated susceptible adults or teenage people. The infection caused by this virus is generally considered severe and self-resolving even in the absence of treatment but in some cases, it needs special treatment and Medicare related to pain and skin itching.

This viral disease as primary infection causes chickenpox (varicella), is characterized by a generalized vesicular rash in children [2]. Virus antibodies in serum and skin lesions of affected children, react with some of the structural proteins of varicella virus where viral immune complex has been demonstrated in sera of patients .The VZV virus can establish latency in the host sensory nerve ganglia, and can reactivate at a later time to cause shingles (zoster) in adults with some problems in immunity, after years or decades [3]. This rate manifested in 30% of usual onset in varicella infections and has been named to varicella zoster (VZV) [3]. Mild chickenpox has been shown to be a rare vaccine associated adverse event, and the vaccine virus strain has also capacity for reactivating and causing mild shingles [4, 5]. In addition, break-through wild-type VZV infections have been shown to occur occasionally in individuals vaccinated previously [6, 7]. The latent infection after reactivation comes with unilateral vesicular rash, ophthalmic, neurologic, viremia, and dermatologic complications. The risk of transmission from zoster to varicella is far lower than from varicella. Varicella develops in people with no history of varicella disease or vaccines.

Serious and painful complications are known to occur regarding chickenpox and shingles. Although both infections are rarely fatal, during the first trimester in pregnancy, varicella can result in congenital malformations in the fetus or disseminated varicella in the neonate depending on the time of infection. There is an effective vaccine for VZV employed a live attenuated by vOka strain, which was developed in Japan in the mid-1970s [3, 8]. This vaccine was licensed for use from 1995 [3].

In the present study, Vero, MRC-5, GPEFC (Primary Guinea Pig Embryo Fibroblast Culture) were used for isolation and characterization of a local varicella virus from suspected varicella patients. A locally authenticated foreskin derived cell line (RFSC) [9] was also introduced as a novel cell substrate for VZV virus production and used for the first time as compared against MRC-5. The virus was cloned after serial dilution by plaque assay. The size of the plaque did not change between 4th and 27th passages. In the upper passages, the yield of virus increased and revealed a higher titer. The presence of varicella virus replication was proved in immunoflurocent using IE62 antibody, and verified through SDS –PAGE, western blotting. The isolated VZV virus was adapted to in-vitro culture in two different cell substrates, during successive passages. Characterization of the virus genome was evaluated in RFLP-PCR using 2 different ORFs (38 and 54). We investigated on, comparative Potency by CCID50 and Real Time after serial passages in Vero cell line. The harvested virus fluid was used for immunological evaluation in guinea pigs. Several VZV challenge trials were also carried out in guinea pigs. The effect of vaccination on mortality and viral load was evaluated. The efficacy of virus in vaccinating guinea pig after subcutaneous (SC) and intra muscular (ID) as compared with intraperitoneal (IP) administration, as well as the humoral and cellular immunogenicity were tested. These immunogenicity results have already shown good results. In this article, we will mention the processes applied and the results regarding isolation and verification, plus identification of a local candidate for the varicella vaccine.

### Ethical consent

The research was conducted in accordance with the Declaration of Helsinki (as revised in 2008) and according to local guidelines and laws. The project was approved by the ethics committee of Faculty of Medical Sciences, Tarbiat Modares University Tehran, and I.R. IRAN. The animal used for the experiment was based on The US Public Health Service’s “Policy on Human Care and Use of Laboratory Animals,” and “Guide for the Care and Use of Laboratory Animals.

## Results

### Sample collection

Varicella virus was isolated from skin lesion in 4 samples. There were different samples showing a positive result for varicella antibodies. Meanwhile, four isolated viruses from all of the samples revealed better result in ELISA test than others, which were selected for primary inoculation in cell culture. The results of IgM and IgG antibodies to varicella virus in serum of these 4 patients are shown in Table 1 and documented as a part of local isolated history. This article is about isolation and characterization of sample number four belong an 8-year-old boy which was negative in mycoplasma testing and after inoculation, it showed better viral lesions in the cell culture.

**Table 1.** Summary of serological results of 4 serum samples which the related vesicular lesions used to isolate the virus in this study.

| No | Sampling date (mo/day/yr.) | Sex | Age | IgG |  | IgM |  |
| --- | --- | --- | --- | --- | --- | --- | --- |
|  |  |  |  | LL<sup>a</sup> | UL<sup>b</sup> | LL | UL |
| 1 | 9/19/2015 | F | 8 | 12 | 20.7 | 0.6 | 1.0 |
| 2 | 8/28/2016 | F | 7 | 12 | 17.6 | 0.3 | 1.9 |
| 3 | 4/10/2017 | F | 9 | 9.2 | 26.1 | 1.9 | 2.7 |
| 4* | 9/6/2018 | M | 8 | 11 | 22.6 | 2.4 | 4.2 |

A: Lower Limit
B: Upper Limit
\* Isolation of virus was successfully performed from sample number 4, belong an 8 years' boys.

### Specific PCR to differentiate local VZV from other Herpesviridae members

After inoculation of the vesicular swab sample/VTM on cell culture by designing the specific pair primer and cloning a conserve sequence in pBHA vector as control positive in PCR test for differentiation the isolate from other member of the Herpesviridae family and the identification of isolated virus as Varicella zoster was performed [1]. In Table 2, the primer set sequence and thermal cycle, as well as the conserve sequence are shown as positive controls.

**Table 2.**
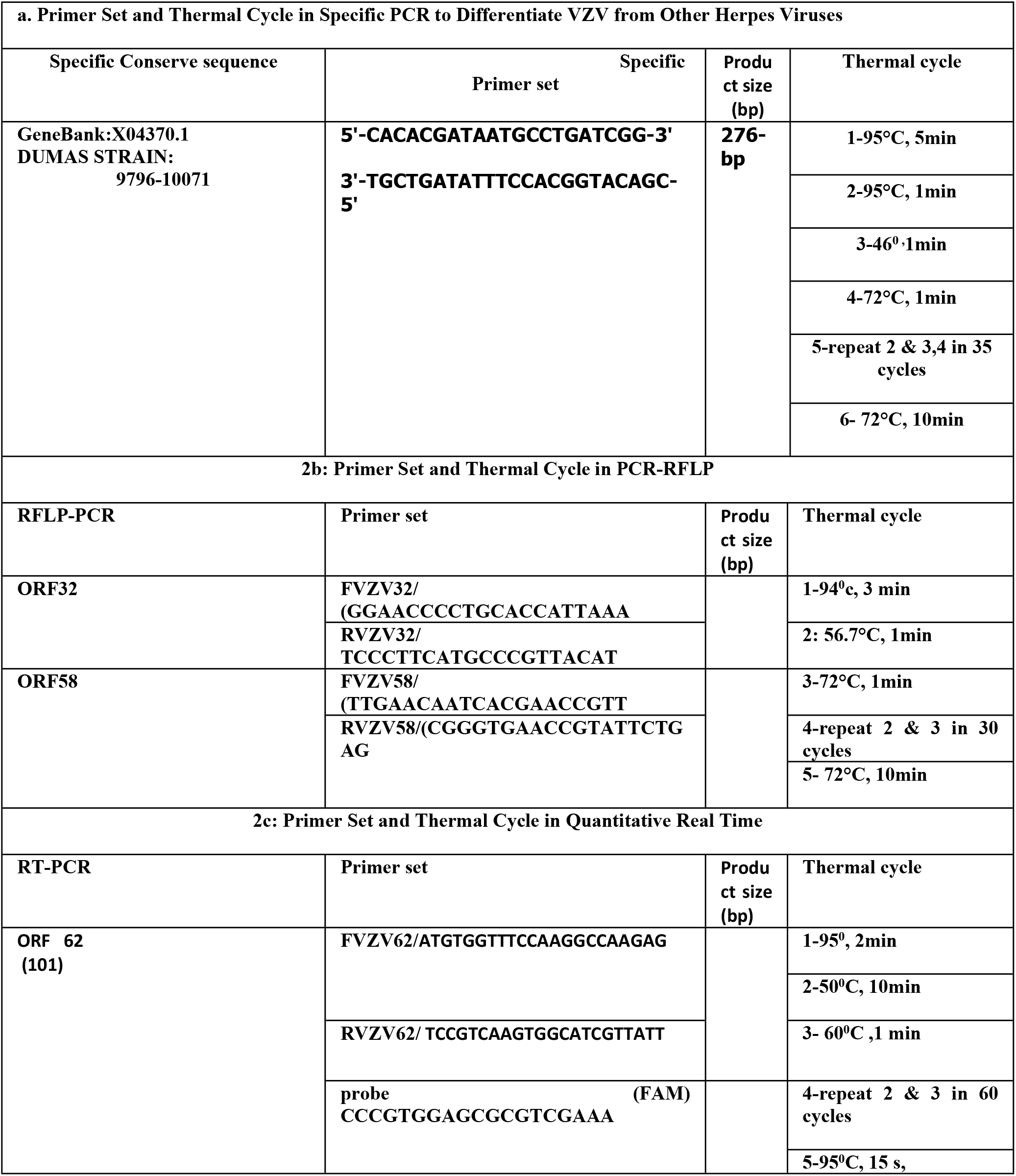
Primer set and thermal cycle in specific PCR to differentiate VZV from other herpes viruses.

The result of a specific PCR test for differentiation of the isolated sample from other Herpesviridae member has been shown 276 bp after electrophoresis on 1.5 % agarose gel, and staining by safe stain. This result was confirmed in comparison with the size of construct as control positive in test.

In Fig 1 the pBHA vector, its primer set and conserve sequence of DUMAS strain cloned as positive control and gel electrophoresis of sample are given the same size.

**Fig 1.**
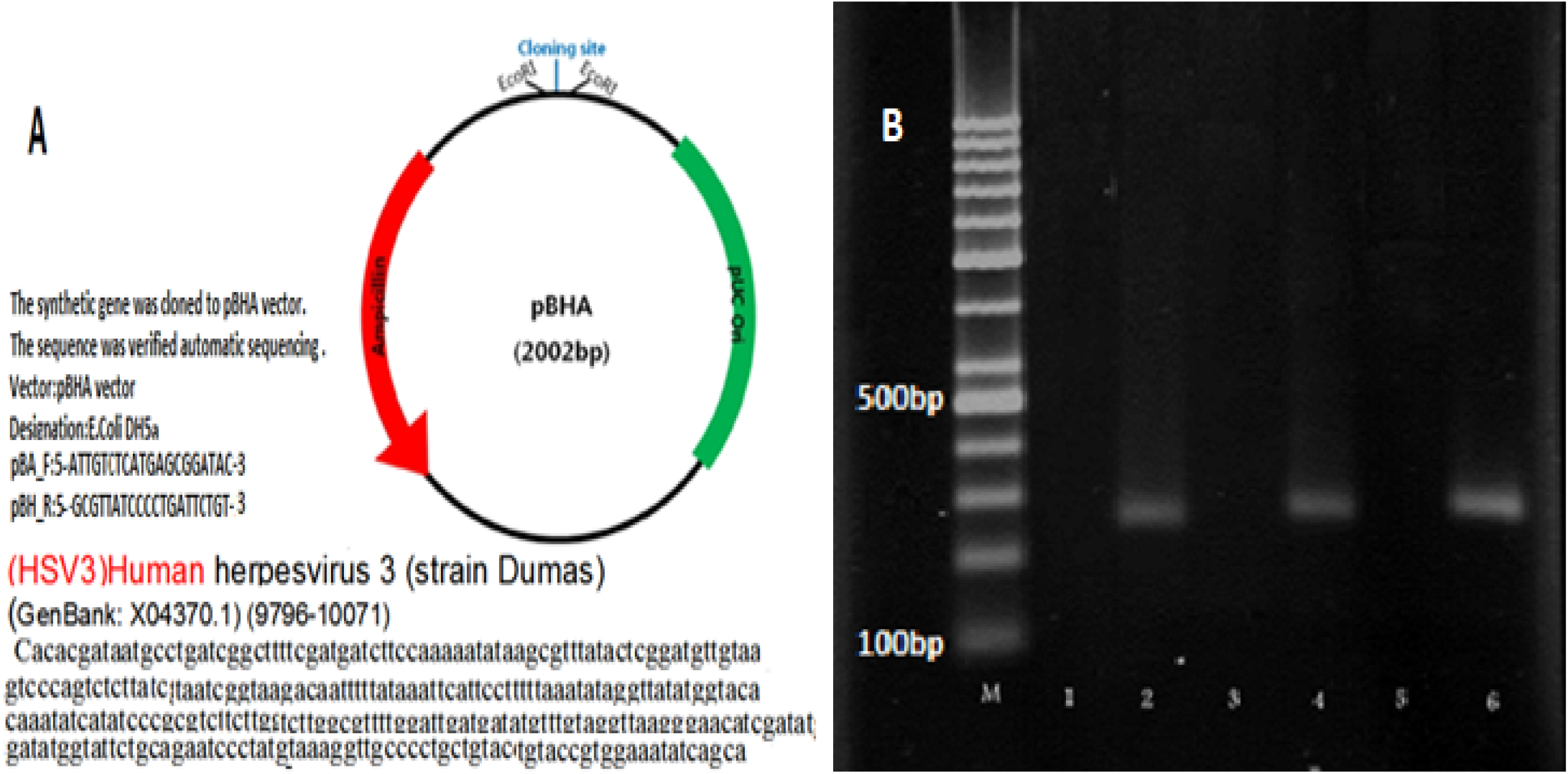
Differentiation of local VZV DNA by specific-PCR. A, pBHA vector; B, Analysis PCR amplification: M lane, Ladder; lane 2, PCR products from control positive (DUMAS Strain: 9796-10071) VZV DNA in dilution 1/10; lane 4, PCR products from local VZV DNA in dilution 1/10; lane 6, PCR products from local VZV DNA in dilution 1/100.

After observing a dense layer of cell line, the transfer medium contain swab in used as sample and inoculated to cell culture monolayer. The culture left in room temperature for 2 hours regarding adsorption of virus. Then, DMEM was added to culture and incubated at 37°C. After 6-7 days, when the cytopathic effects (CPE) appeared on culture, the flask was frozen and thawed. The cells and the flask supernatant were centrifuged at 3000 RPM for 20 minutes. The supernatant and pellet were extracted by Qiagen DNA extraction kit and underwent the extraction by Qiagen DNA extraction kit according to the manufacturer’s user manual. The DNA was amplified by specific PCR. At each step, using specific primer pair of varicella zoster with appropriate concentration. DNA detected in agarose gel using ethidium bromide and comparison with markers to verify the presence or absence of the virus. Positive control was tested along with positive infectious sample. Comparison of the results showed that the components were similar to the positive control.

### Virus isolation and attenuation

CPE in cell culture systems is not always clear enough on primary inoculation, thus tissue culture fluids may have to be passaged at least 2-3 times for full adaptation and more than 20 times for isolation and attenuation as we did such process for VZV virus from swab sample in this research. To follow the microbial and viral adventitious agent, for every 5 passages, a sample from isolated virus was inoculated in three different cell lines. The mycoplasma and sterility testing were done based on requirement [5]. Two days post inoculation the cultures were examined microscopically to monitor the virus growth. The culture media changed with fresh ones every 2 days. Once the virus cytopathic effect spreads on the culture, the media were harvested and stored at −60°C for next step. Serial sub cultivation was carried out and continued more than 25 times to reduce virus virulence and adapted in growth at 33 ° C. The first eleven passages were on the MRC-5 cells, then the cell substrate was changed by primary guinea pig embryo fibroblasts cells (GPEFC) (using 20-day pregnant fetus) and viruses were passaged from 12 to 24 on this cell substrate. In the third stage, MRC-5 cells were used again and virus culture continued until passages number 30. In parallel, all the processes were performed with the local foreskin cell line, as an alternative to MRC-5 for future production [10]. For this purpose, the growth kinetics of two cells in terms of higher titer production of isolated virus were also compared (Fig 2).

**Fig 2.**
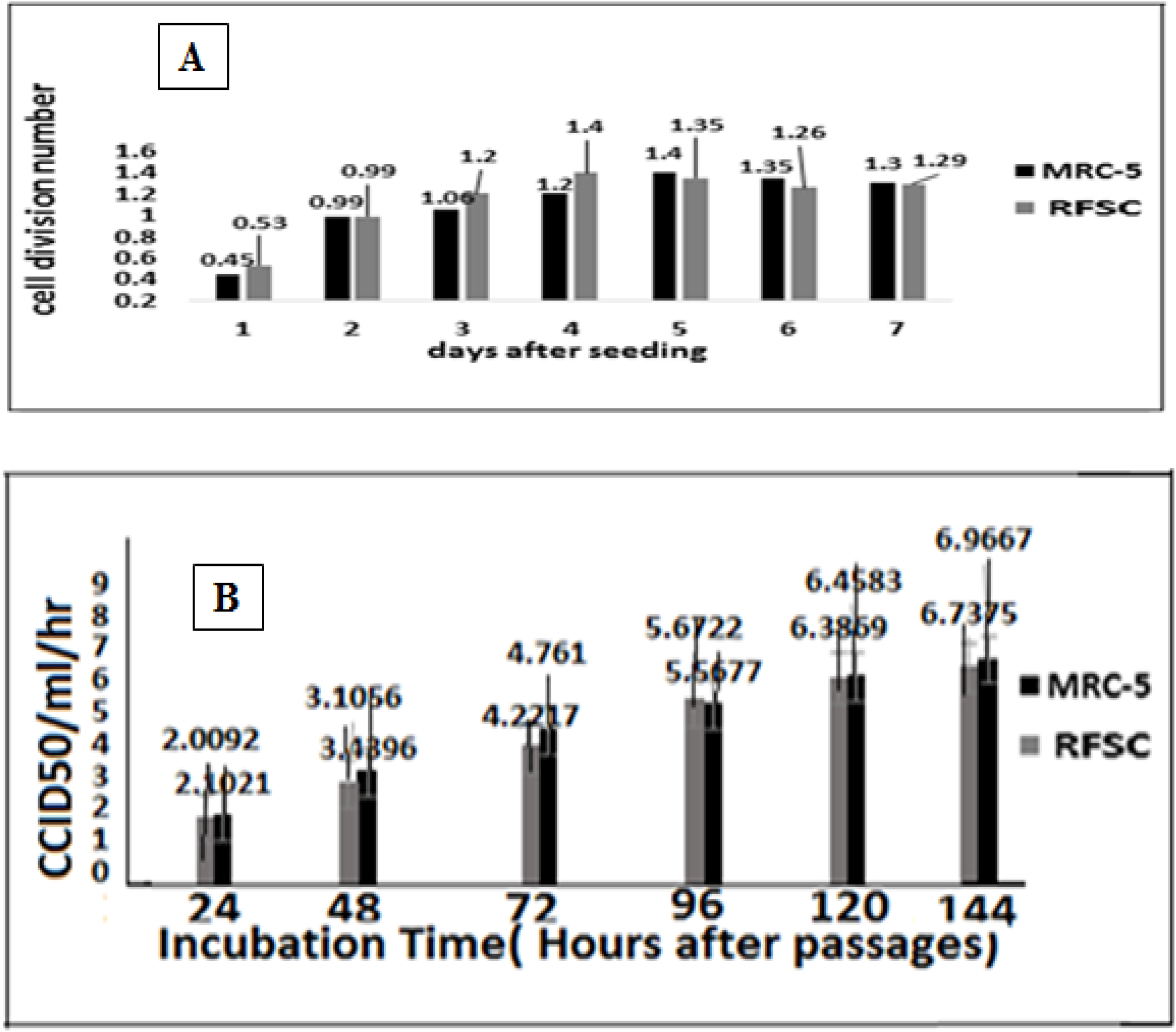
VZV on human diploid lung and human foreskin cells. (A) Comparative cell division number between two human cell substrate during seven days after seeding. MRC-5 as standard human diploid cell strain and RFSC as a continuous local human foreskin cell substrate: The local RFSC at passage 65 and MRC-5 at passages 26. (B) Replication characteristics of local VZV isolate in human diploid lung (MRC-5) and human foreskin (RFSC) cells: Two human cell lines were inoculated with the local VZV and its CPE assessed by CCID_50_ method. Replication kinetics of isolated VZV were compared with RFSC as an alternative cell substrate for future research and production.

### Comparative evaluation of local virus replication on two different cell substrates MRC-5 and RFSC

In the present study, concerning the limitation to access to MRC-5 causes, RFSC, a local cell substrate that developed based on requirement by Razi Vaccine and Serum Research Institute from the foreskin, was used to produce VZV vaccine using the local virus.

In the present experiment, the replication kinetics of the two different passages viruses were compared on MRC-5 and RFSC cells infected at two MOI: 0.01 and MOI:0.1. Local isolated VZV showed peak virus titers by day post-infection, for high and low passages post infection (Table 3).

**Table 3.**
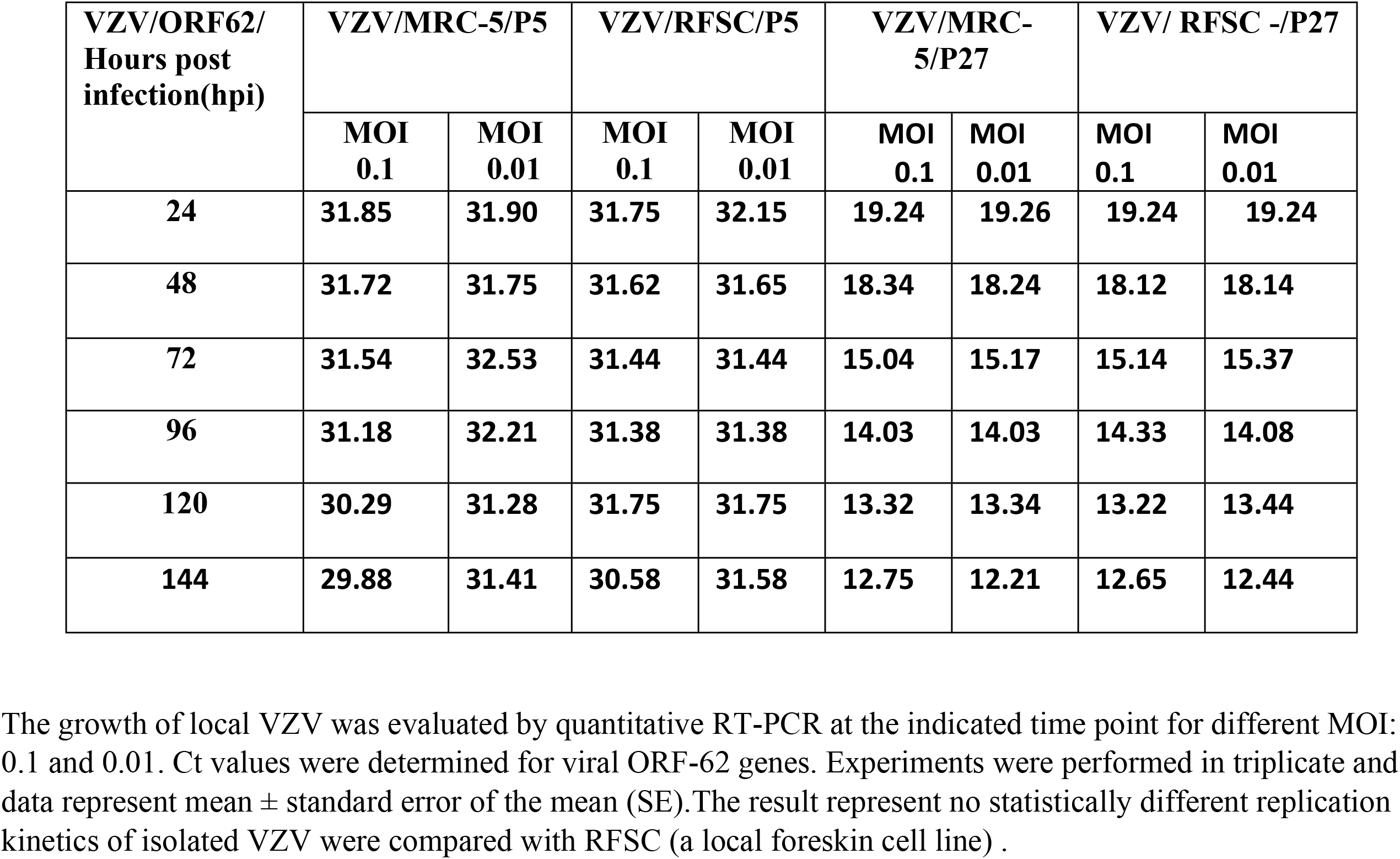
Viral ORF-62 genes expression levels on MRC-5 and RFSC cell substrate.

| VZV/ORF62/<br>Hours post<br>infection(hpi) | VZV/MRC-5/P5 |  | VZV/RFSC/P5 |  | VZV/MRC-5/P27 |  | VZV/ RFSC -/P27 |  |
| --- | --- | --- | --- | --- | --- | --- | --- | --- |
|  | MOI<br>0.1 | MOI<br>0.01 | MOI<br>0.1 | MOI<br>0.01 | MOI<br>0.1 | MOI<br>0.01 | MOI<br>0.1 | MOI<br>0.01 |
| 24 | 31.85 | 31.90 | 31.75 | 32.15 | 19.24 | 19.26 | 19.24 | 19.24 |
| 48 | 31.72 | 31.75 | 31.62 | 31.65 | 18.34 | 18.24 | 18.12 | 18.14 |
| 72 | 31.54 | 32.53 | 31.44 | 31.44 | 15.04 | 15.17 | 15.14 | 15.37 |
| 96 | 31.18 | 32.21 | 31.38 | 31.38 | 14.03 | 14.03 | 14.33 | 14.08 |
| 120 | 30.29 | 31.28 | 31.75 | 31.75 | 13.32 | 13.34 | 13.22 | 13.44 |
| 144 | 29.88 | 31.41 | 30.58 | 31.58 | 12.75 | 12.21 | 12.65 | 12.44 |

The growth of local VZV was evaluated by quantitative RT-PCR at the indicated time point for different MOI: 0.1 and 0.01. Ct values were determined for viral ORF-62 genes. Experiments were performed in triplicate and data represent mean ± standard error of the mean (SE). The result represent no statistically different replication kinetics of isolated VZV were compared with RFSC (a local foreskin cell line) .

Infection with each of the three (high and low passages local isolated beside vOKa) viruses proved to be typical syncytium-cytopathic in these cells, which remained persistently infected for the 10 days follow-up period, though virus titers started to increase from 3 DPI onwards. In addition, peak virus titers were measured by 9-10 DPI in MRC-5 and RFSC cells for high and low passages viruses and by 7 DPI for vOKa as control positive. The result showed similar titers were obtained from MRC-5 and RFSC cells inoculated by low passages (5^th^) compared to strong higher titers obtained from P: 17, P: 21, P: 27, infected MRC-5 cells. To study the short-term dynamics of virus replication, a one-step growth curve was performed on MRC-5 (human lung diploid cell strain) at an MOI of 0.1. The eclipse period for all stage high passages viruses was between 16 and 24 hours on MRC-5 cells. Slightly different eclipse periods were found on human foreskin cells (between 18 and 30 hours), but it was not significant compared to MRC-5. At 7 DPI, 60% of the cells were inoculated with MOI of 0.1 showing the cytopathological effect and high titers detected in the culture supernatant of both MRC-5 and RFSC cells.

In a comparative evaluation of MRC-5 and RFSC cell substrate, although the growth of the virus has increased, it was not statistically significant. Viability and recovery rates of stored cells were assessed, in both RFSC and MRC-5 cell categories, where survival rates were 94% and 92%, respectively, which is same as the levels we expected since both cell lines had the same characteristics regarding the growth kinetics(Fig 3).

**Fig 3.**
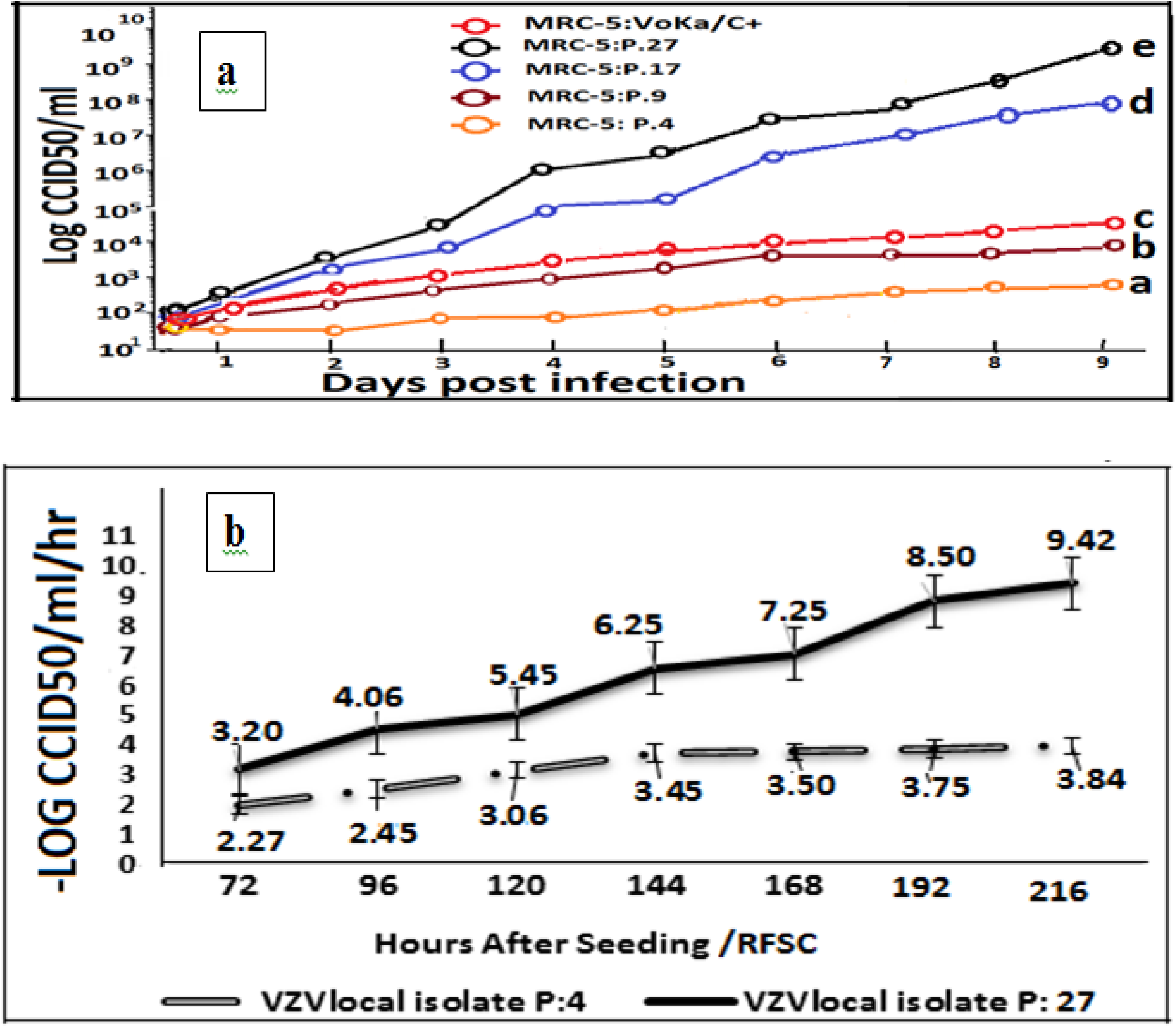
Replication and potency assay of Local isolated VZV virus during different passages on MRC-5. (a) Development of VZV virus inoculated on MRC-5 cell substrate. CCID_50_/mL of VZV virus titers were measured by potency tests as described in the text. The values of, higher and lower level of virus titer was assessed from; p27. (8.75-9.40 LogCCID_50_) and P4. (2.75-3.40 LogCCID_50_). This level for Control positive was (vOKa3.7-4.3LogCCID_50_) after 9 days’ post inoculation. The virus at P.27 and P17 (6.5-7.4 LogCCID_50_) was significant in compare of P9. (3.40-4.5 LogCCID_50_) and P4. (2.75-3.40 LogCCID_50_). Increasing the particles number to 6 and 5LogCCID_50_ for p 27 and P17 was linear to level of virus in RT-PCR but was significant. (b) Replication and potency assay of local isolated VZV during different passages of VZV on local cell substrate (RFSC). After successive serial passages, the virus has become adapted to growing in new cell substrate. This procedure increases the proliferation and number of virus particle. The virus titer has increased by about 6 log CCID_50_/ml in high passages (p: 27), compare to low passages (p: 4) and produces more particle in potency testing at hours post inoculations daily and also after 9 days in high passages The titer of the virus was measured using Karber method.

To evaluate the effect of the adaptation on virus production, we performed potency assay in the cell culture. We harvested fluid containing isolated virus from MRC-5 and RFSC and inoculated on Vero cells through logarithmic serial dilution. We obtained the result of CCID_50_ by the Karber method in Vero cell lines (Fig 4). We used Vero cells as it is highly sensitive to quality control of the harvested virus. Successive passages showed the complete adaptation of RFSC to virus as well as to MRC-5. It was observed that after 3 consecutive passages, the complete adaptation of the virus to RFSC cell was established. Regarding MRC-5 on first passages, the cytopathic effect of isolated local VZV and control positive was visible after 8-9 days post-infection. Although viral cytopathies were seen in the first passage, the rate of virus adaptation to the cell substrate was not high enough to produce the high yield varicella zoster for future vaccine production, and several passages were given to the isolated virus to accustom to the in vitro culture.

**Fig 4.**
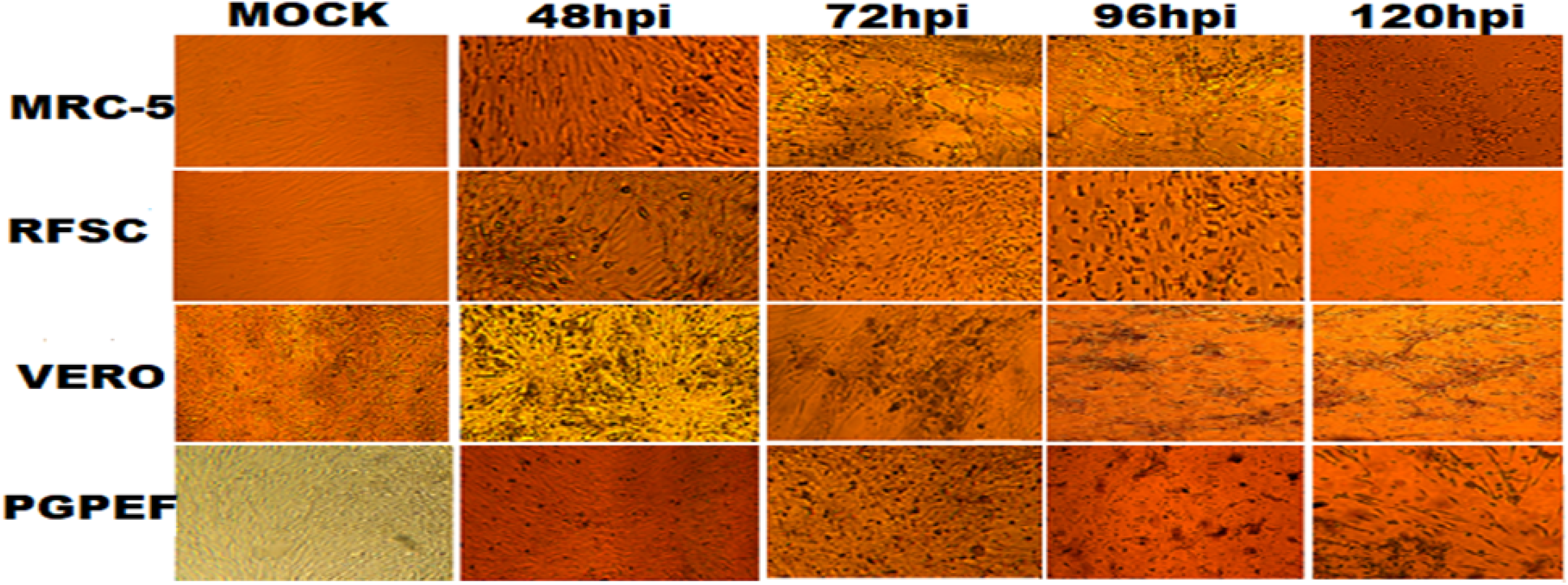
The local VZV cytopathogenicity. Human lung diploid cell (MRC-5) and human foreskin fibroblast (RFSC), Vero, and primary guinea pig fibroblast cells (PGPFC) to local VZV virus infection at 48 and 72and 96 and 120 hours after infection (100x magnification).The virus inoculate as MOI: 0.1 and was in 17th passage.

In this procedure, vOKa was used as control positive. As the passage of the virus increases on the target cells, so does the number of viral particles. The number of particles in passage 27 as the high shows a huge difference with the low passage 4, The virus does not need adaptation to the MRC-5 and its growth was achieved after first passages by sub-culturing of the virus during 8-9 days. Local VZV virus presented typical CPE and caused RFSC-cell lysis in a monolayer, which were easily observed through microscope visualization. The pH of the media for the virus culture was optimized on 6.6±0.2. The total cell per 25cm^2^ flask, were 4.2×10^6^ ± 1.5×10^5^ as same as for MRC-5. These optimized growth conditions for local VZV on RFSC cell substrate caused achieving a higher yield of virus in potency. Maximum virus productivity of the local VZV virus was high in RFSC (10 ^9.42^ CCID_50_/mL).

### Plaque Assay

The most appropriate method for isolating a single particle from a virus in a sample from different genetic populations is the plaque method. Using this method, through measuring the plaque size in some viruses, an attenuated from virulent parent of a virus can be indicated and differentiated. This phenomenon has been reported for many viruses after induction of serial passaging in cultures [6, 11].

In the plaque test, the fresh monolayer of Vero cell was infected with a serial dilution (10^**6**^-10^**12**^) at MOI equal to 0.1 from local VZV virus. The culture incubated at room temperature allowed the virus to be absorbed into the cells and then covered with a nutrient medium containing agar. In this procedure, virus-infected cells released viral particles, which limit the spread of new viruses only to adjacent cells. As a result, infection with any viral particle creates only one area of infected cells that can be seen with the naked eye called plaques. The plaque test using neutral red as vital dyes help see the boundary between the plaque and the surrounding single-layer cells, thereby absorbing the dye in living cells. Thus, the bright plaques resulting from the virus replicating in a field of red, and the living or non-living infected cells are distinguished. Since a virus particle is sufficient to form a plaque, the virus titer can be presented as a plaque-forming unit (PFU) per milliliter. In addition to titration, we used the plaque method to purify and clone the virus. In the present study, the above method was used to evaluate the titer and clone the isolated virus (after 25^th^ passage) and also to evaluate the size of the plaque in relation to its attenuation. We determined plaque sizes of the passaged viruses. We did not detect any changes of the virus phenotype. Plaque areas were measured using electronic size counter (Fig 5).

**Fig 5.**
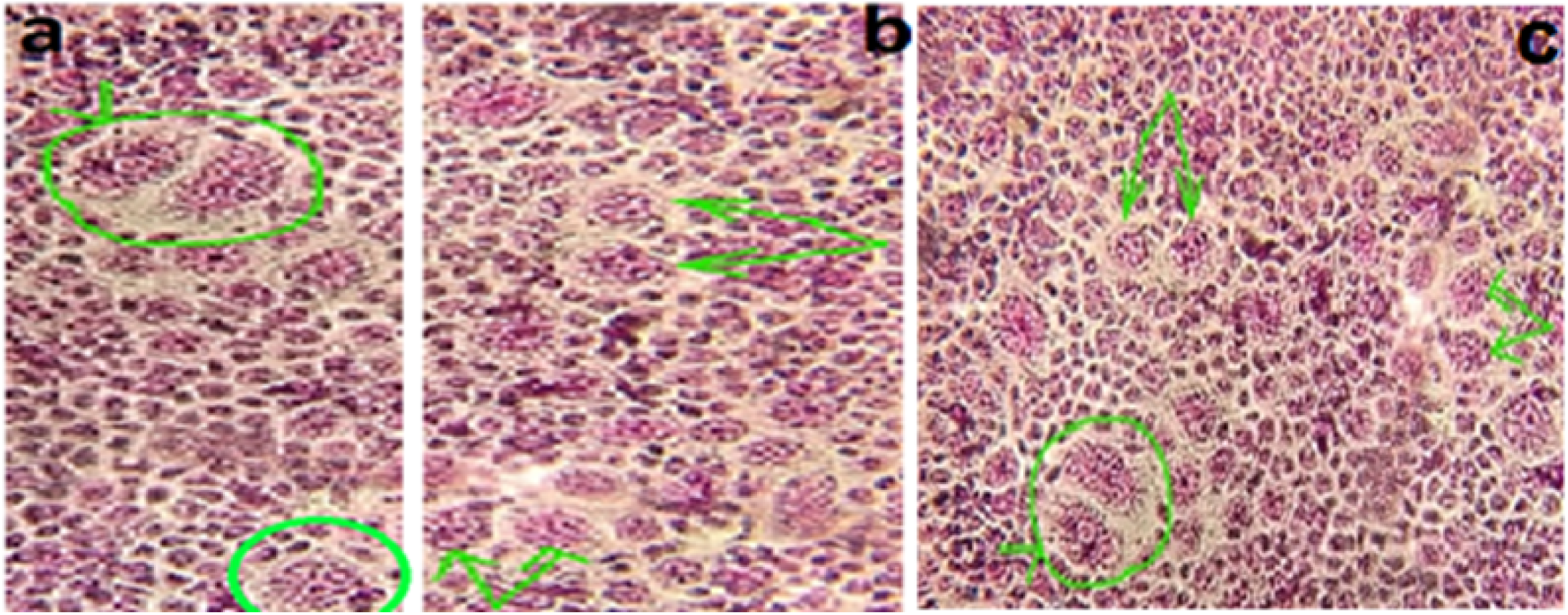
Plaques formed by the viruses that were passaged in cell culture (Scale bar, 200 μ). (a) Representative Images of plaques formed by viruses that were passaged 5 times serially at low multiplicity of infection in cell culture. (b) Representative Images of plaques formed by viruses that were passaged 17 times serially at low multiplicity of infection in cell culture. (c) Representative Images of plaques formed by viruses that were passaged 27 times serially at low multiplicity of infection in cell culture. Serial passaging of viruses has not altered phenotype of plaque of viruses. Although the plaque produced in the 27th passages of virus were larger, and plaques formed by the 4^th^ passages of virus were smaller. But the size distribution of relative plaque diameter normalized against the average plaque diameter of the parental virus. P-values were calculated using multiple comparison test, indicates P<0.05.

### Characterization of isolated VZV viruses by RFLP-PCR after different passages in tissue culture

The isolated viruses were examined by PCR and restriction fragment length polymorphism (RFLP) analysis, using the ORF38 and ORF54 region. Initially, we performed PCR by designing two pairs of specific primers for these two ORFs (Table 3). Then, via digestion through two restriction enzymes, BglI and PstI with a fully -optimized thermal cycle, digestion was performed on the PCR product. In electrophoresis of PCR and RFLP products on 1.5% agarose, the results were confirmed after safe staining (Fig 6).

**Fig 6.**
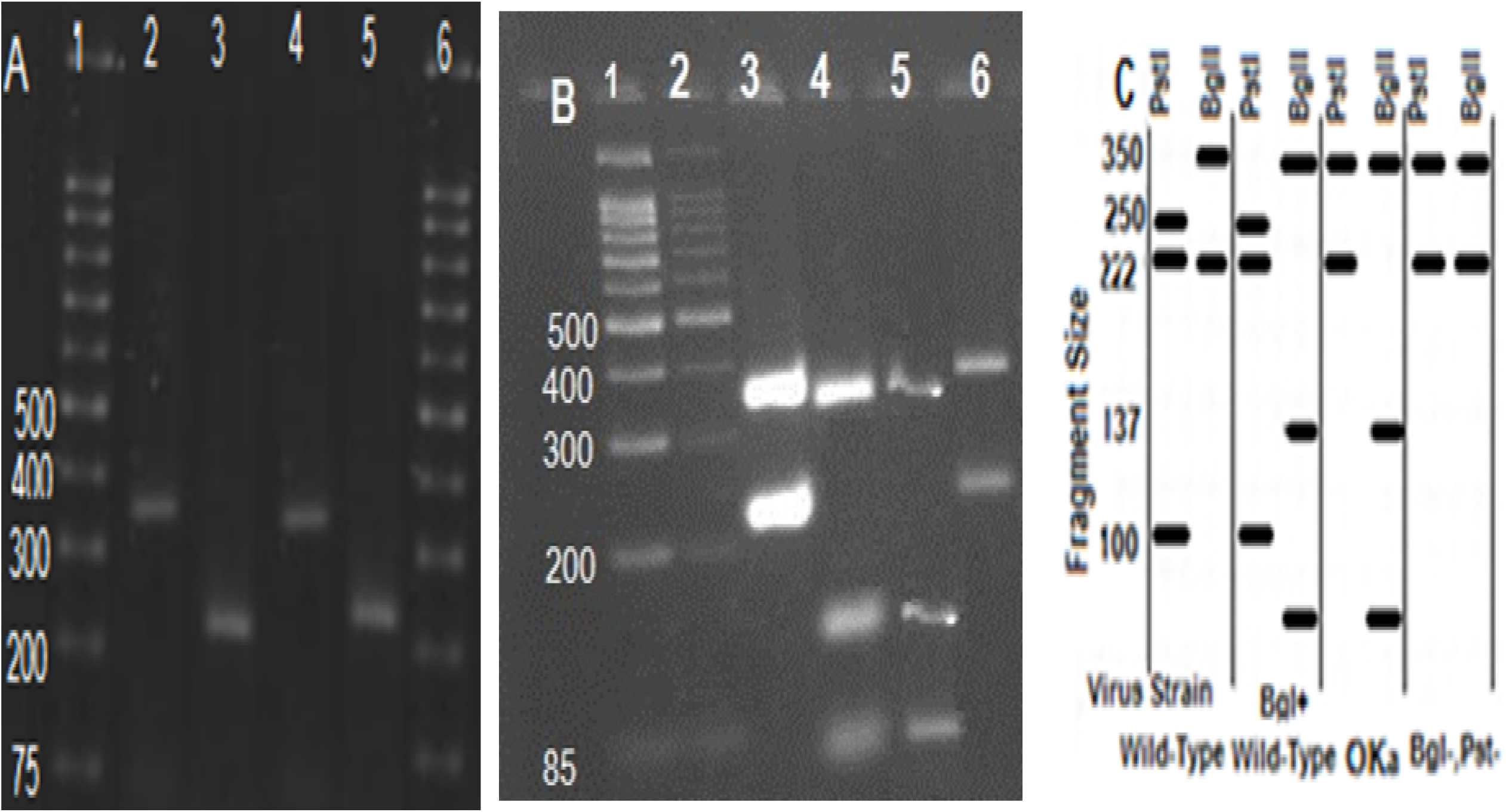
Results of electrophoresis following RFLP test on PCR products of high passage local VZV DNA by RFLP-PCR. (A) PCR amplification of ORF 34 and ORF58, contain 222-bp and 350-bp fragmentlane1, ladder; lane 2and 3, PCR products from vOKa as reference positive, 350-bp and 222-bp; lane 4and5, PCR products from isolated varicella, 350-bp and 222-bp. (B) RFLP on PCR product to analysis the fragment size differentiates wild from attenuated isolated; lane1 and 2 ladder; lane 3, vOKa profile after digestion by PstI (350-bp and 222-bp); lane 4, vOKa profile after digestion by BglI as reference positive (85,100,137bp fragment size); lane 5, digested products from local isolated VZV by BglI (85,100,137bp fragment size); lane 6, digested products from isolated varicella (222 and 350-bp fragment). (C) Schematic RFLP fragment size BglI and PstI.

### SDS-PAGE Western blotting analysis

The activation of ORF62 was evaluated by Western blotting following SDS-PAGE. In this study, a band was identified related to IE62 protein in varicella virus. The protein is the most important viral activator of VZV gene transcription (Fig 7).

**Fig 7.**
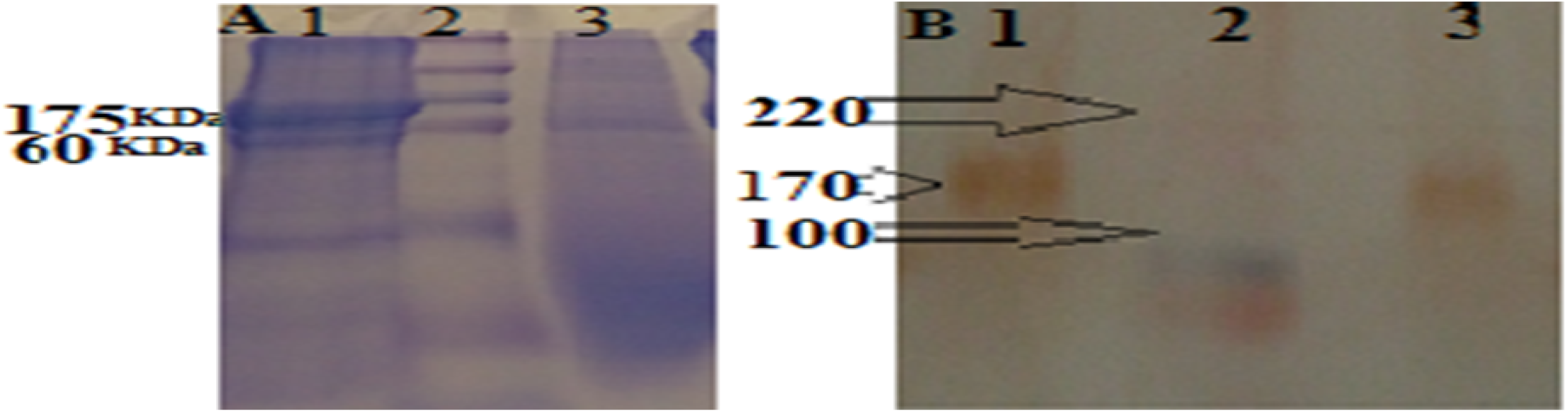
Comparative SDS-PAGE and Western blotting results between isolated VZV and vOKa as control positive. (A) SDS-PAGE of isolated VZV and control positive; Lane 1, 175 kDa band of IE62 protein in the local VZV; Lane 2, Protein Prestain marker; Lane 3, 175 kDa band of IE62 protein in positive control. (B) Western blotting of isolated VZV and control positive; Lane 1, 175 kDa band of IE62 protein confirmed with specific antibody in the local VZV; Lane 2, Protein Prestain marker; Lane 3, 175 kDa band IE62 protein confirmed with specific antibody in positive control. This result confirmed identification of IE62-specific protein as the most important viral transcription activator of VZV gene between isolated VZV and vOKa as control positive. Its presence was examined using labeled specific antibodies, in the Western blot experiment with color band observation. Binding of the labeled secondary antibody was associated with the anti-C-terminal monoclonal antibody near the C-terminal IE62 protein, along with the identification of positive control.

### Study on expression of the local isolated VZV gene by specific antibody (anti-IE62) in IF test

Our goal was to determine whether cold adaptation is an applicable strategy for attenuation of local isolated VZV herpes viruses. To study the effect of growth condition (viral replication) and its optimization at a lower temperature (viral attenuation), we recorded the IE62, as DNA Pol is essential for virus replication in vitro and in vivo, and we expected that reduction or increase of the effective protein on its function and production might change the biological properties of the isolated viruses, which could be easily observable both in vitro and in vivo. As observed previously the replication of high passages increases, if the conditions for the growth of the virus in 300C are optimized. This feature markedly presented in potency testing following that, but the number of fluorescent particles in IE62 antibody testing showed a sharp decrease in P:30 compared with parental P:5 because of occurring probably attenuation. The reduction of expression of IE62, following sub culturing of the virus, from low to high passage for cell-substrate MRC-5 in IE62 antibody testing by IF has been shown in Fig 8.

**Fig 8.**
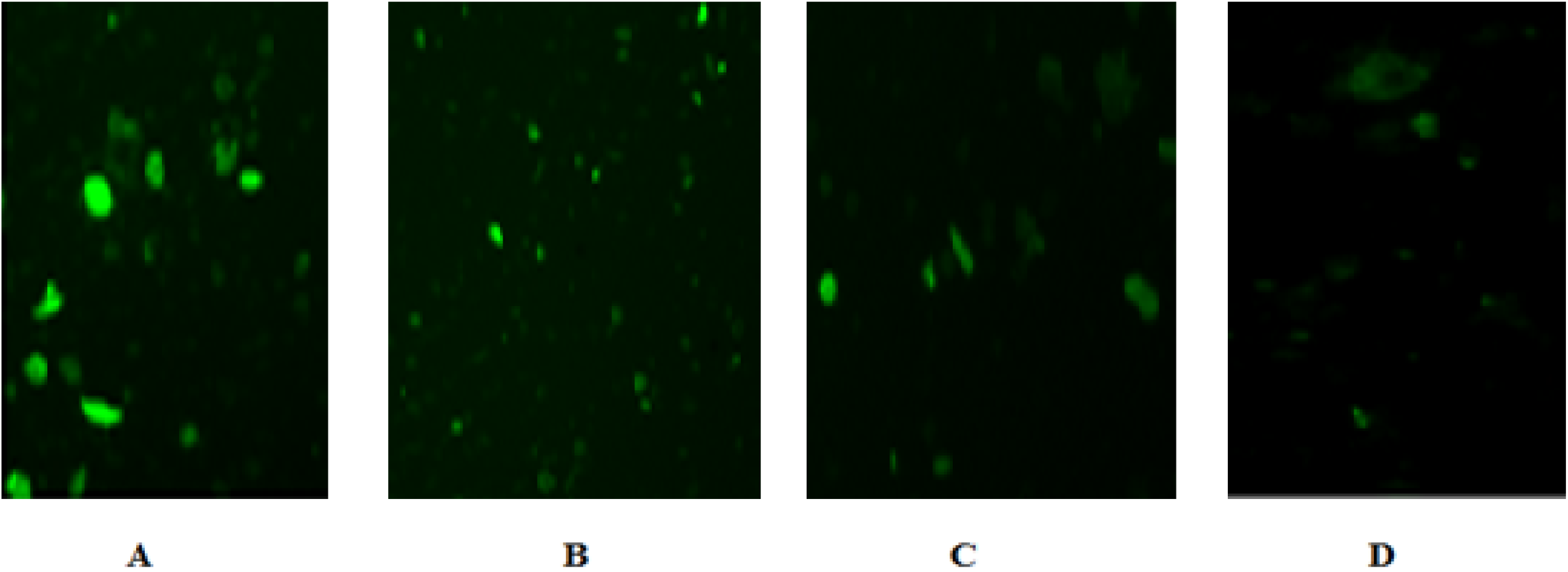
The local VZV in IF test after serial passages. A, the 5th passage of local VZV; B, the 30^th^ passage of local VZV; C, control positive virus; D, the uninfected cell.

### The comparative replication rate of local isolate VZV virus by CCID_50_ and RT-PCR (Real Time PCR)

The replication rate of local isolate VZV virus in human cells was evaluated by quantitative RT-PCR for 0.1 and 0.01 MOI.

The kinetic replication of ORF62 gene in infected cells was similar to the virus titration results. VZV infection induced a rapid increase in the levels of ORF62 gene in MRC-5 cells. There were not any significant differences in the transcription levels of the gene in VZV-infected cells. In this test, the levels of viral ORF62 genes from infected cells were analyzed by one-step quantitative RT-PCR. The reactions were conducted in duplicate and carried out for 30 cycles. The relative expression threshold cycle (Ct) value was normalized using internal control. Reactions were repeated three times for each sample. Representative images in Fig 3 show MRC-5-transfected with a culture fluid at MOI: 0.1. To quantify virus production was driven by different passages, the harvested fluid was inactivated and we analyzed DNA production from the parental and high passages 72 h after cell culture transfection. qPCR analysis showed that 27th passages well adapted and produced significantly more as compared with the 4th passage(Fig 9). This difference was completely significant. The purpose of the above test was to evaluate the rate of ORF62 gene replication, which reported that the increase in gene proliferation could be attributed to limit reduction.

**Fig 9.**
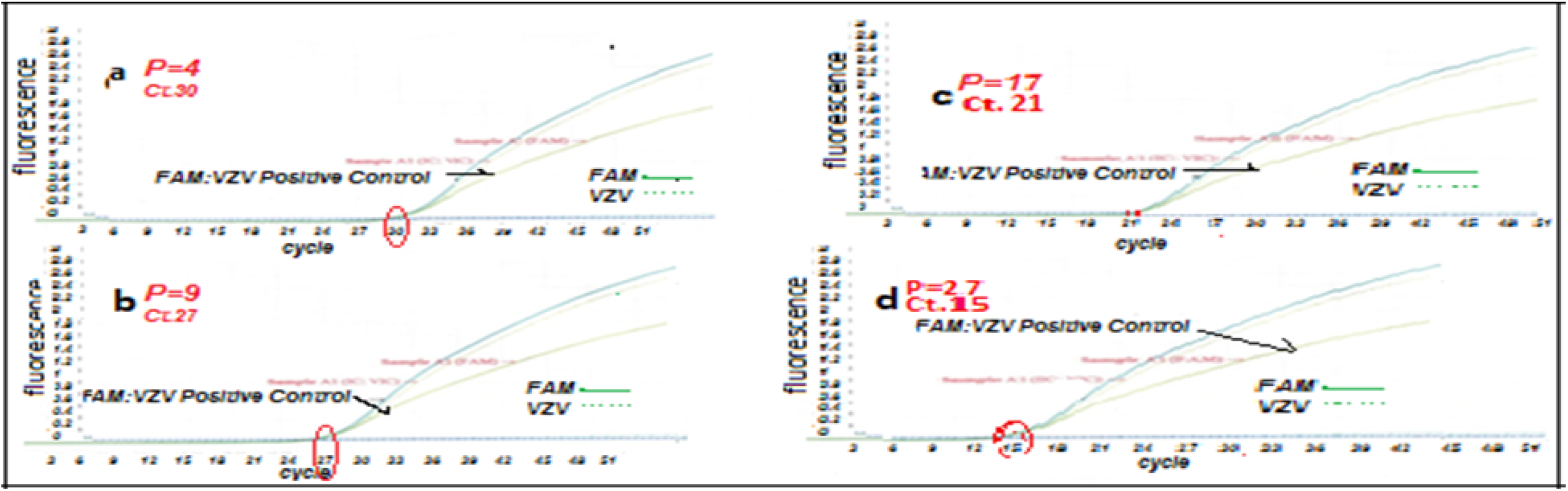
Viral particles in current VZV isolate from Iran by RT-PCR on MRC-5 (72hpi). (a) Isolated VZV-infected MRC-5 cells P.4. (b) Isolated VZV-infected MRC-5 cells P.9. (c) Isolated-infected MRC-5 cells in P.17. (d) Isolated-infected MRC-5 cells P.27.

### Immunological studies and serum antibody development

This procedure was done to evaluate the immunogenicity of local isolated virus and its efficiency for producing the antibody as dose finding.

Several studies have demonstrated the serological and pathogenesis of VZV virus. We sought to determine the kinetics and optimization of minimum dose of local isolated VZV that can induce antibody production. We found that isolated VZV was serologically positive in the animal model inoculated with 10^5.0^ CCID_50_/mL (group A), 10^4.3^ CCID_50_/mL (group B), 10^3.3^ CTCID_50_/mL (group D), and 10^3.0^ CCID_50_/mL (group E), either ID or SC. In the animals as early as 6 days post inoculation, no clinical signs, fever, redness, and weight loss were apparent from 1st days post-inoculation onward. There is a significant increase in antibody titters between the first stages of blood sampling belonging to pre-inoculation of the virus and the second stage which is 6th days following the first inoculation. Similarly, blood samples were taken on 21^th^ and 44^th^ and 162^th^ days after the virus was injected an ascending trend is also evident in almost all groups in their antibody titters. More animals that received 10^4.7^ CCID_50_ /dose, 10^4^ CCID_50_ /dose 10^3^ CCID_50_ /dose or 10^2.7^ CCID_50_ /dose from high passages (30^th^) developed IgG antibody compared to animals receiving 10^2.7^ CCID_50_/dose low passages (5^th^: group F). As shown in Fig 10, an increase in Ig M was observed for up to 21 days. Then, a gradual decline from 44^**th**^days to 166^**th**^days after injection was recorded. An increase in Ig G was clearly seen after 44 days and progressed to 162 days post-injection (Fig 11). Note that group H includes animals exposed to vaccinated group A without any injection, to assess the transmission of isolated virus. A case of death or disease symptoms, and an increase in VZV not observed in the last group. The result of serum antibody development revealed the differences between high and low passaged viruses.

**Fig 10.**
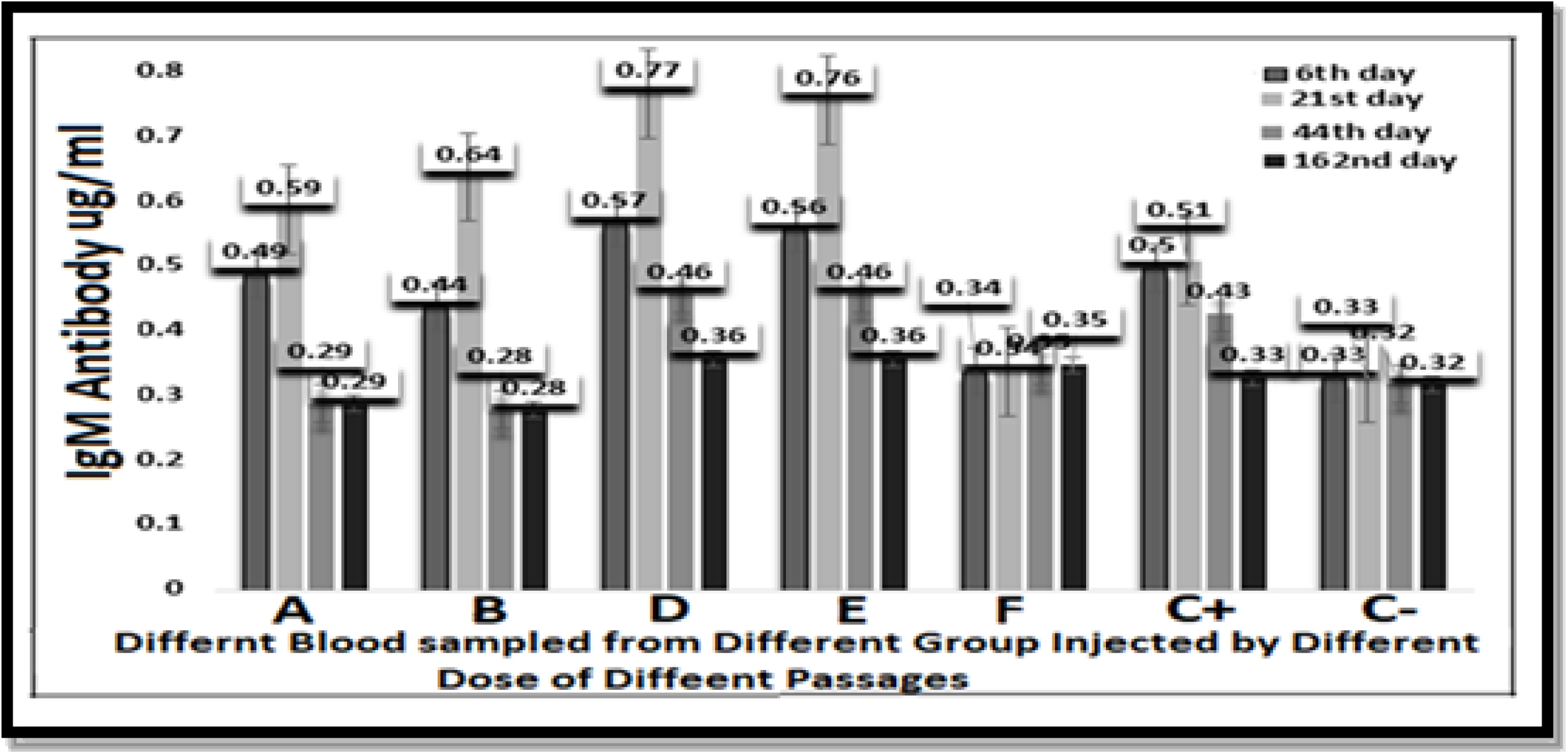
Development of local VZV-specific IgM over time in (350-405 gr) Guinea Pig. Guinea Pig were inoculated ID or SC with (A) 10^5.0^CCID_50_, (B) 104.3 CCID_50_, (D) 10^3.3^ CCID_50_ and (E) 10^3.0^ CCID_50_ of high (30th) passages isolated VZV virus, (F)10^3^ CCID_50_ of low passages (5th) and (C+) 10^3.3^ CCID50 of vOka vaccine as control. The lines indicate threshold level of the assay (P>0.05).

**Fig 11.**
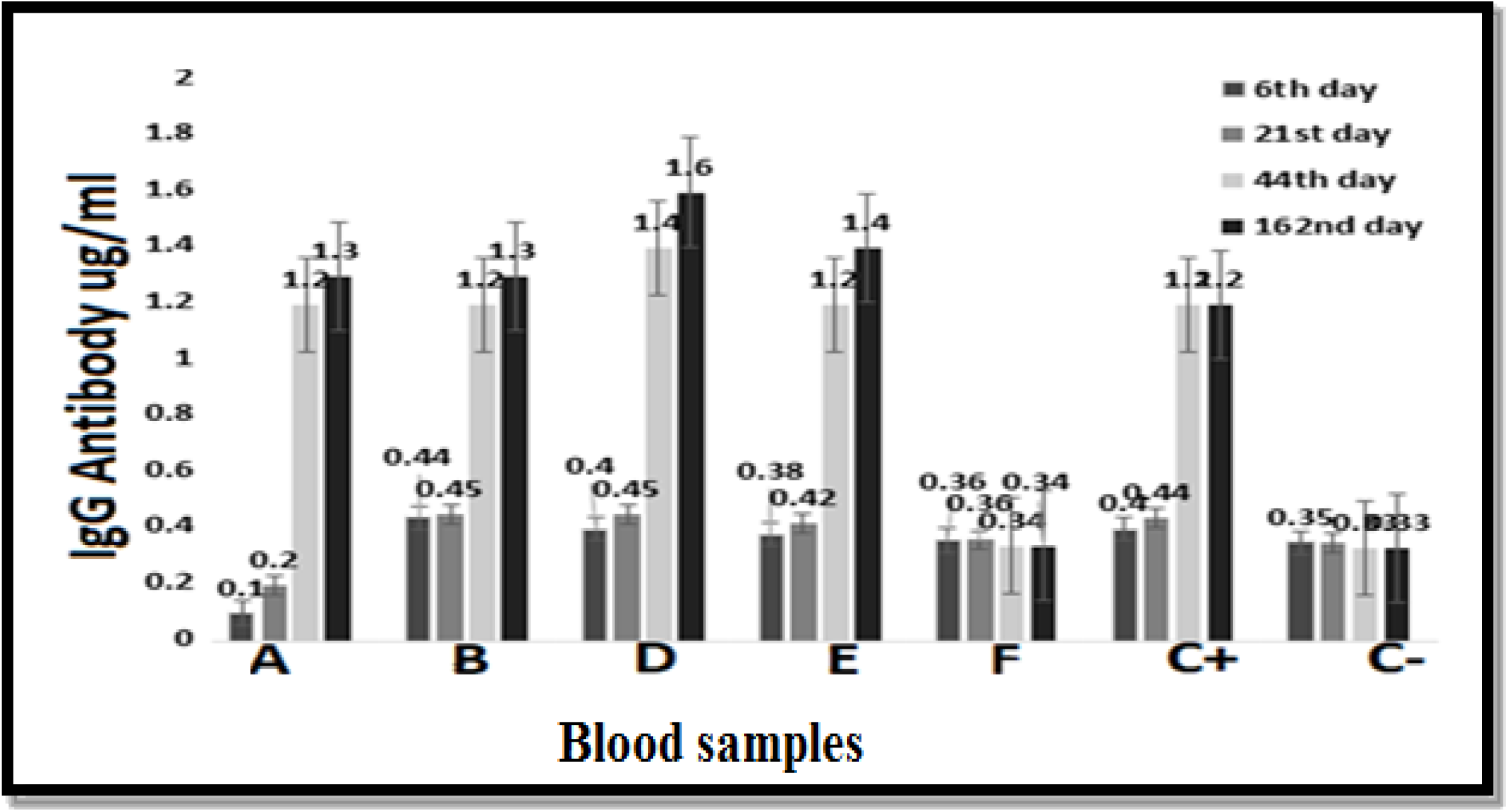
Development of local VZV-specific IgG over time in Guinea Pig (350-405 gr). Guinea Pig were inoculated ID or SC with (A) 10^5.0^CCID_50_, (B) 104.3 CCID_50_, (D) 10^3.3^ CCID_50_ and (E) 10^3.0^ CCID_50_ of 30th) passages isolated VZV virus, (F)10^3^ CCID_50_ of 5th passage and (C+) 10^3.3^ CCID50 of vOKa vaccine as control. The lines indicate threshold level of the assay (P>0.05).

The virus did not antibody in the serum samples of animals inoculated with 10^3^ CCID_50_/mL _**of**_ low passages (5^th^). We compared antibody production in animals inoculated SC with high as attenuated and low as wild isolated virus and vOka commercial vaccine (group C^+^) as positive controls. All animals had to be alive until 9 months after inoculation and monitored weekly. The development of antibodies was not detected in negative group animals (group C^−^) which had received water for injection (WFI). The route of inoculation did not influence significantly the antibody production of animals receiving any dose of virus at 6, 21, 44, or 162 days post-inoculation (P>0.05). The use of isolated virus as antigen in the viral neutralization (VN) could explain the neutralizing antibody titres that high passages isolated virus (P: 30) inoculated guinea pig developed after inoculation with 10^3.3^CCID_50_ or 10^3.0^ CCID_50_ (Fig 12) [12].

**Fig 12.**
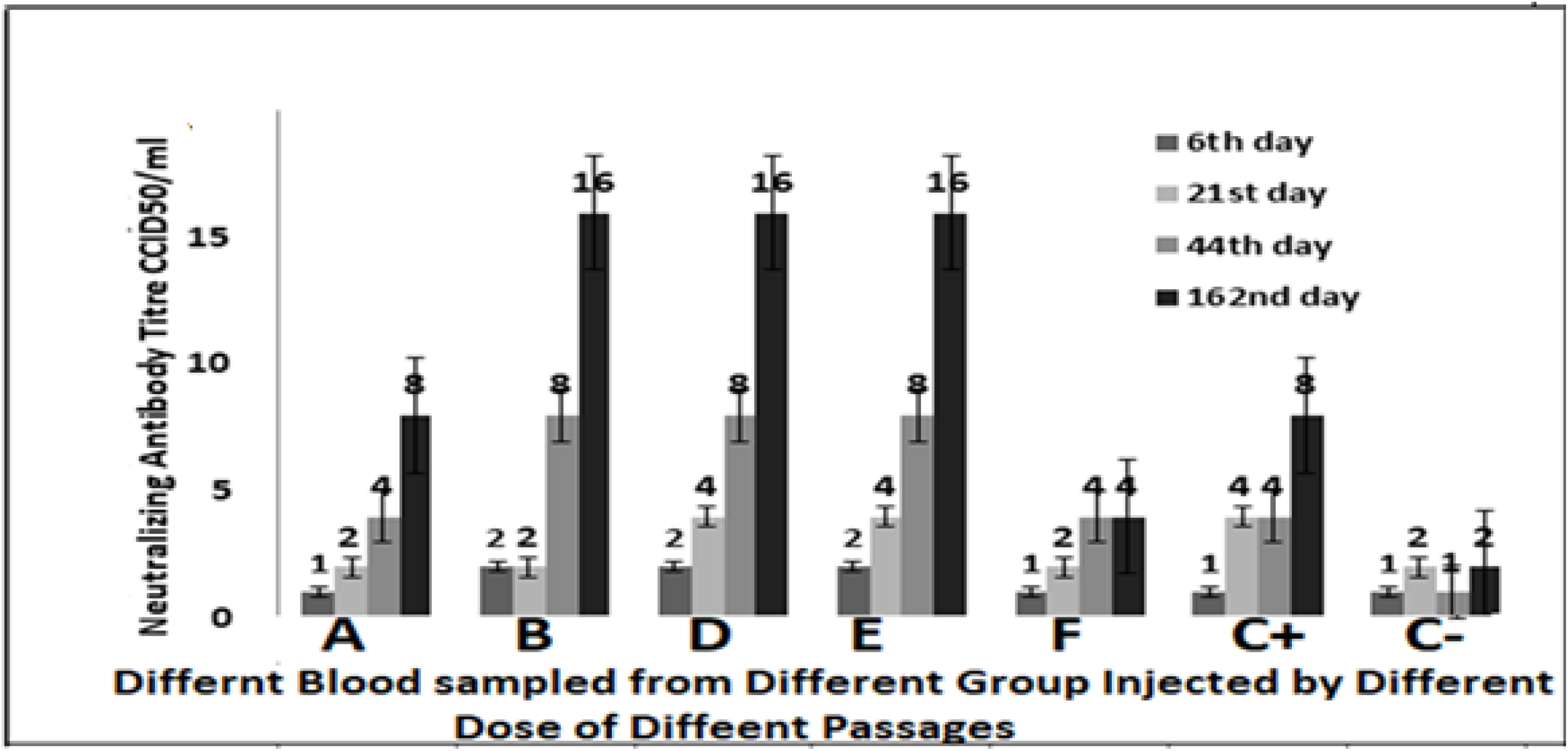
Development of Local VZV-specific neutralizing antibodies over time in (350-405 gr) Guinea Pig. Guinea Pig were inoculated ID or with (A) 10^5.0^CCID_50_, (B) 10^4.3^ CCID_50,_ (D) 10^3.3^ CCID_50_ and (E) 10^3.0^ CCID_50_ of high (30^th^) passages isolated VZV virus, (F)10^3^ CCID_50_ of low passages (5^th^) and (C^+^) 10^3.3^ CCID_50_ of vOKa vaccine as control. The lines indicate threshold level of the assay.

### Skin test

On the other hand, stimulation of cellular immunity following receiving isolated VZV virus was shown in test animals, on the 21st day using ID injection of positive control virus suspension, after 24-72 hours with a skin reaction in the form of redness and stiffness more than 5 mm in diameter, in the injection site.

## Discussion

The live attenuated varicella vaccine using the vOka strain is the only vaccine licensed to prevent human herpes virus infection. Immunity and Safety to this vaccine has been demonstrated in early clinical trials in Japan [13]. Clinical evidence for attenuation of vOka has shown that vaccine-related rashes occur in only about 5% of healthy children and adults. Proliferation of the vOka VZV virus during immunization of immunocompromised individuals or those using immunosuppressive agents is relatively limited [14, 15].

Lower yields of infectious virus and slowness of skin lesions associated with decreased viral protein synthesis of skin xenografts in SCIDhu mice compared to pOka proposed that the attenuation of vOka vaccines results from a reduced capacity to replicate in skin, which allows a longer interval for the adaptive immune response after viral inoculation. According to this hypothesis, infectious vOka may be cleared before skin lesions formed and before cell-associated viremia develops [16, 17].

There are different experiences in which surveyed succeeded to isolate the viral agent from the swab lesion, sera, CSF of affected patients by cultivation of sample in different cell line. In some of them the isolated virus remained highly potent and went under following passage for future vaccine [6, 18].

In this article, via a similar technique, we describe the in vitro isolation of local varicella zoster from an 8-years boy skin vesicle swab. This sample selected from four other because not only blood serum of patient presented acceptable antibody level but also presented good condition regarding sterility, mycoplasma, cell culture, as well as its subsequent molecular and biological characterization. The differentiation from other Herpesviridae family member was 100% identical to the primary isolate as varicella zoster by specific PCR assay. Chickenpox infection is highly contagious from days 7-21 of the commune (average 14 days), and studies suggest that the VZV particle is transmitted through the air. Airborne particles from skin lesions are the main source of infection and are more involved in the spread of the disease than the patient’s respiratory secretions [19, 20]. The risk of transmitting chickenpox is about five times higher than in those with shingles [21, 22].

The disease begins with symptoms such as lethargy and fever from one to two days before and after the rash. The fever usually lasts for 5-7 days in parallel with the rash. Each rash appears as an itchy lesion within 24 hours of the stage from macular to papule and then to the vesicle and then the pustule and eventually to the wound. Thus, all of these lesions are seen in a person with chickenpox, where mainly the trunk, scalp, and even epithelium of the internal organs (endothelium) also become infected [23, 24]. In the case of VZV, the phenomenon of escape from the immune system as well as escape from binding to antibodies has been observed, which leads to asymptomatic recurrences [12]. The virus invasion might not be dependent on the amount of virus replication. Given the nature of transmission via usually close contacts superficial scratches or respiratory secretion, in the human infection, it is generally believed that the VZV virus is invasive at low doses [25, 26]. The amount of virus that has been mentioned for human infection in various articles is closely related to the immune status of the elderly or other terrestrial diseases. Indeed, varicella zoster virus causes infection in a substantial number of adults or infants. Further comparisons have revealed that either route of infection or the age of the case led to marked differences in the clinical outcome, antibody response, or amount of viral RNA recovered from the patients’ infectious material. Although the incidence of varicella disease is declining as a result of vaccination but there is not including as expanded program for immunization in Iran yet. In order to study the replication potential of the isolated virus, different experiments were performed in this research. The shortage and limitation to access to MRC-5cell strain caused to candidate an authenticated local cell line from foreskin, as alternative for future research and development [10]. The growth curve of VZV could reveal no significant differences in replication kinetics between the titre in different passages of viruses, either on skin epithelial or human fibroblast cells. The mechanisms underlying these differences are not completely clear. It is possible that the two different human cell lines differ in viral receptor density or qualitative or quantitative differences in anti-viral response to infection. Alternatively, differences in the rate of cell division or split level in culture could also explain the differences observed in the yield virus potential of the two human cell substrates. On the other hand, RFSC is a continuous cell line and MRC-5 is a diploid cell strain capable of limiting the number of passages for production. In this regard RFSC, can be cultivated continuously and will have a higher value in terms of production. The results revealed that the increase in virus yield in these two human cell substrates: MRC-5 and RFSC was due to the isolated virus adapted to cultivation at 30°C. Our data confirm the high sensitivity of the two different human cell lines after inoculation by different passages of local isolated VZV virus.

One of the factors that can determine the rate of adaptation to the cell is the rate at which the cells become infected with the virus. Initially, the virus is attached to the cell surface, and then the number (percentage) of cells that the virus has been able to spread is determined as virus replication. A higher adaptation rate is equal to the higher virus growth and ultimately the virus yield. The specific rate of virus production in cell culture is actually the rate at which the virus proliferates in the cell over a specific period of time before the cell was lysis or disrupted. The higher the rate of virus replication, the greater the rate of virus production in the culture and the shorter the time required in terms of high yield virus production. To examine the process of replication and viral yield in a cell line, determining the rate of potency efficiency will be one of the important factors for vaccine production. The higher this factor, the higher the viral particles in a given volume of cell substrate, which is very important from the point of view of vaccine production.

Herpes viruses, as many other large DNA viruses, are equipped with their own DNA replicase, which renders them independent of the host DNA replication machinery. The catalytic subunit of the herpesvirus DNA polymerase is a pivotal enzyme responsible for genome replication, and thus, for successful transmission of genetic information from one generation to the next [4, 7, 27].

IE62 protein is the most important Trans activator for VZV DNA polymerase. Some studies have suggested that the level of expression of IE62 protein inversely correlates with virulence. However, more recent data suggest that IE62 protein expression levels are not critical for pathogenicity [28]. It remains to be seen if the IE62 protein of VZV would have the capacity to induce such signals and confer virulent or attenuated phenotype. IF test showed serial passages at low temperature associated with decreasing IE62 expression in high passage of the local isolated VZV, in contrast low as wild isolated virus. Thus, as other investigations on vOka have suggested one of the characteristic signs of VZV attenuation can be reduction of IE62 function, it was investigated by other researchers [29].

The virus has had more than 30 passages on both cells in parallel. Virus stocks were prepared by 30 additional passages of the primary isolate on human cells (MRC-5, RFSC) for each 5 passages interval. The purpose of this phase was to induce in vitro adaptation of viral isolate at lower temperatures than what would occur in the body during natural infection. Due to successive passages, not only has the virus become accustomed to growing at lower temperatures, but the result has seen an increase in the titer of the virus in higher passages than in lower ones. Remember that different amounts of defective interfering particles in the harvested viral fluid may have affected the replication kinetics of viruses. The result of such an adaptation in many articles as a method that reduces virulence and produced attenuated virus vaccine strains has been reported [4, 30].

To study the cytopathogenicity of human cells to local VZV virus, MRC-5, RFSC, GPFE and Vero cells were infected at MOI: 0.1. Nevertheless, the marked CPEs were evident 48 hpi for MRC-5 cells infected at targeted MOI, which increased with time. At 72 hpi, approximately half of the infected cells appeared smaller and irregularly shaped compared to mock cells (Fig 4). Infection of MRC-5, RFSC, and also GPFE cells by local VZV virus caused a fast CPE and a rapid fall in the pH of cell in culture. The granular and fragmented cells became obvious within 48 to 72 hpi. In contrast, infected Vero cells showed the rate increased slightly from 24 to 48 hpi. Note that higher viral titers were observed in infected MRC-5 cells, and the peak of titer reached 124 hpi. Thus MRC-5 indicates more sensitivity for VZV proliferation. This is important in terms of optimizing the best time for cell passages and virus inoculation. The isolated virus passaged more than 25 times in MRC-5 cell strain, regarding reduction in virulence. Also, subsequent experiments were carried out with (P: 5) as the parent isolate (P: 5) that had been passaged fifth on MRC-5 and (P: 30) that we hope was attenuated. The results mainly showed a significant difference between primary characterized isolate (P: 5) and the current passaged virus (P: 30) virus in term of potency and RT-PCR.

One feature of the serial sub culturing in human diploid fibroblast that differed from the others was the adaptation of the strain to growth at 300C which we included in our procedure. The idea behind this was to rapidly induce attenuation, on the basis of previous experience with attenuated measles, mumps and rubella strains [11, 31–33]. By following the work of others showing that replication at low temperatures selected against virulence. By the 27th passage in a human diploid cell strain, we expected local VZV had clearly become attenuated for humans.

In the follow up by potency testing in CCID50, it was found that the infectivity titer of the isolated VZV became higher as the passaging in cell substrate progressed. This feature was clearly confirmed in the RT-PCR test results (Fig 9). In other words, we found a correlation between virus levels determined by CCID50 /mL and mRNA levels determined by qPCR. The differences between primary characterized isolate (P: 4) virus mainly showed significant in results of potency and RT-PCR of the high passaged virus (P: 27). Consistently, viruses or RNA could be detected in inoculated cell culture in-vitro and laboratory animals in vivo. Similarly, in the present article the virus was isolated from infectious material such as vesicular swab and throat, especially in 3-4 first days of infection from VZV patient case. We could determine the virus titer in the infectious material (blood serum) of the patient by endpoint dilution. The quantitative PCR data suggest that the sample used for detection of patient was enough containing approximately 10 TCID50/ml of infectious virus. These results are in agreement with results from plaque assay, which showed that the number of plaques increased. Since decreasing the virus growth temperature is one of the factors influencing the reduction of the virulence, we have accustomed the isolated VZV, to growth at 300C. We expected that, increasing the virus titre at a temperature lower than normal in human body (300C) should result in phenotypic changes, which would be easily observable in cell culture. Fortunately, these results were confirmed by three different quantitative tests. Increasing the virus titre determined by CCID50 /between higher (27th) and lower passage (4th) and increasing the Ct in FAM-q PCR related to ORF 62. Although the above results show that the increase in passage has led to an increase in virus titre, the size of the plaque has not been affected. The plaque size between different passages resulted serial passaging of viruses has not altered phenotype of plaque in local isolated VZV viruses. Although at a glance the plaque produced in the 27th passages of virus were larger, and plaques formed by the 4th passages of virus were smaller in size, the size distribution of relative plaque diameter normalized at 27th against the average plaque diameter of the parental (4th) virus. In calculating the test the P-values resulted P<0.05. This marker characteristic was applied by many of other researchers regarding the attenuation of different isolated virus [11, 30].

As prior in vivo study of the virus proved to be necessary to further study the characteristics of the virus, we studied on the local isolate VZV by PCR-RFLP, after more than 25 passages. The restriction fragment length polymorphism analysis (RFLP) on DNA is recommended as one of the most valuable molecular techniques that can distinguish the complete analysis between different VZV strains [8, 34, and 35]. The isolated strain in tropical regions such as Africa, Bangladesh, China, India, Central America and northern Australia also reported as BglI+ strains. The result of many studies also reported RFLP profiles of wild type VZV strains in Japanese isolates as, PstI+, BglI+ or PstI−, BglI+ as in US, UK, Europe PstI+, BglI+ or PstI−, BglI+, meanwhile in eastern Australia result of varicella RFLP-PCR showed PstI+, BglI− .Using this method, RFLP profile of isolated varicella in this article, after more than 20passages showed PstI^−^, BglI^**+**^(PstI site and BglI site in many Asian strains) [36–39].

Our findings are in agreement with previous studies. RFLP-PCR of the local isolate proved to be highly identical at nucleotide level to the other isolates from Asia for which the complete genome sequences are available and published. The identity between this isolate and others belonging to Asia, Europe, African and other region isolates is necessary to repeat the experiment and for accurate analysis [39, 40].The main limitation of this article was the lack of complete sequence result of from local isolate presented in this article and other isolates from different provinces in the country. If we could examine other isolations from other provinces in terms of complete sequence, we could be more certain about the isolation of the country’s circulation. These limitations were due to financial and logistical reasons. These processes by excess financial support is under second repeat for isolated virus in this article. Phylogenetic analysis of viral particle has been largely based on the complete genome sequence [41]. At the time of analysis there were some financial limitations to repeat the process for accurate analysis but the problem solved right now the result is under work. Further studies are required to clarify the biological relationship between the differences observed between local isolated in the present paper and its evaluation with various other VZV isolates.

In our studies we did not see differences in antibody development between the different routes of inoculation. In skin test we could not detect **“**redness and inflammations**”** side effect of replicating virus at the site of inoculation early after infection via ID. There was no association between development of seropositivity and route of inoculation. In animals inoculated with 10 ^2.7^ CCID_50_/dose of isolated virus at 5^th^ passages no specific antibody response was detected. In order to assess whether the inoculated virus was sufficient to induce antibody response, in the skin test with positive control virus injection, the safety status of animal models receiving the high dose virus was tested with the result obtained indicating that it was stimulated [42, 43,44]. Comparison of in vitro and in vivo characteristics and virulence after adaptation to growth in 30° C and more than 30 sub passaging with those of vOKa vaccine (commercial vaccine) showed clear similarities with vOKa –VZV strain [13]. In contrast, low passages (p5), act as non-attenuated and exhibited a different tropism for cultured cells, different antibody development, and nearly 100% different from its daughter at 30th passages. Overall, all animals exhibited signs of progression developing neutralizing antibodies by the three different doses of high passages isolated viruses. Extensive development of the antibody in guinea pigs by VZV correlated with the characteristic signs of attenuation. Given the differential antibody development patterns of the three high passages isolated virus with these viruses in low passage, it would be interesting to compare the receptor usage of the parent low passages as wild with high passages daughter as attenuated. These findings as compared with positive control suggest that in immunogenicity, the three high passages viruses are most likely similar to vOka as attenuated commercial vaccine [13]. Our findings are in agreement with previous studies suggesting the used method serial passages and following results.

Both the in vitro and in vivo systems for isolated propagation described in the present study will be valuable tools to further elucidate the isolation and adaptation, as well as the process for developing a candidate for research.

## Materials and Methods

### Sample collection

The samples contained skin lesions belonging to an 8-year-old boy. The swab sample was received in viral transmission medium (VTM) which contained sufficient amounts of antibiotics.

### Sample screening by PCR

To confirm the existence of the varicella and its differentiation from other herpes virus, a pair of primers was designed along with positive control. It was used to test the samples and differentiate it from other Herpes viruses in the sample [, 45, 46, 47].

### Virus isolation and attenuation by monitoring cell infection

Four different cell lines were used for isolation of varicella virus in tissue culture: a) African green monkey cell line (Vero) prepared in tissue culture tube, measuring the potency of isolated virus b) human diploid cells (MRC-5) prepared in tissue culture flasks (25cm2), c) local human foreskin cell line (RFSC), d) Primary Guinea Pig Embryo fibroblast cell culture (GPEFC) grown in tissue culture flasks (25cm2). The growth medium for all cultures was DMEM containing 5-8% of in house irradiated bovine calf serum. All of cultures tested for sterility and mycoplasma and were negative. The cell sheets were washed once with PBS before inoculation. The specimen was inoculated as 0.5 ml into each cell line. The maintenance media was composed of DMEM with 1-2% of in bovine albumin serum added after 2h adsorption at room temperature. The cultures were incubated at 300°C in stationary phase. The isolation methods were designed based on our previous experiment and World Health Organization WHO requirement (Two days’ post inoculation the cultures were examined microscopically to monitor the virus growth. The culture media changed with fresh ones every 2 days. Once the virus cytopathic effect spread on the culture, the media were harvested and stored at −600C for the next step. Serial sub-cultivation was carried out and continued more than 25 times to reduce virus virulence and adapted in growth at 30 0C.

### Viral growth kinetics, specific virus production rate, and overall virus productivity at 30°C by CCID _50_ method

Specific growth characteristics of MRC-5 cell and the yield of the isolated VZV virus in this cell strain were captured as follows. Tripsinated 4×175 Cm2 flask of MRC-5 and counting the cells. Inoculated with MOI: 0.1 from local isolated VZV virus. The culture flask was stirred at room temperature for 15 minutes at 20 rpm, after which a sufficient amount of DMEM containing 5% FCS + 50μg Gentamycin was added as culture medium. This mixture was divided as 2×10^6^ cell / 25cm^2^ flasks (80000 cell / cm2) and incubated at 36 ° C. After 48 h, the culture medium was replaced with a serum-free medium. The flasks were incubated at 32-33 °C. At 24-hour intervals, three flasks were removed from the incubator and sampled from the supernatant to determine the titer of the virus. Then, the flask was Tripsinated and the cells counted the rate of virus production was calculated. The specific rate of virus production in the cell culture was determined by the same method.

The higher the growth rate, the more cells it can produce over a period of time; thus, the faster the product can be produced and the more efficient it can be. The culture fluid was harvested from cell line daily for 240 hours. To assess the comparative adaptation and replication number of viral particles in the cell line, potency assay was performed The 0.1 ml of culture fluid, inoculated in 10-fold dilutions (10^−1^ - 10^−9^) on Vero cell substrate cultured in tube. All dilutions were done in 4 repeats. The viral growth and its cytopathic effect were investigated for 12–14 days using an inverted microscope. The potency of virus was calculated as CCID_50_/mL. This method was done simultaneously for three times in parallel for both passages (4^**th**^and 27^**th**^) in 300c (Fig 2). The commercial (**v**OKa) vaccine strain was used as a positive control.

### Plaque assay for purification of isolated virus

Briefly, MRC-5 (1**×** 10^**5**^) seeded in a well of a 24-well plate was infected with 100 PFU of local isolated virus. The entire inoculation was performed in duplicate. Cells were frozen and thawed 2, 3, 4, 5, 6 and 7 days post infection with 10-fold serial dilutions inoculated onto fresh MRC-5 and Vero. Plaques were stained and counted six days post infection. The plaque on MRC-5 from the highest dilution of the isolated virus was used for cloning. To analyze cell-to-cell spread of viruses’ plaque areas were determined. Briefly, the fluid harvested from MRC-5 containing 50 PFU was mixed with Vero (1×10^5/ml^) and seeded into a well of a 24-well plate. After 6 days, plaques were visualized via indirect immunofluorescence (see below). For each virus, images of 50 randomly selected plaques were taken at 100-fold magnification. Plaque areas were measured using image software from which, assuming that ideal plaques would have a circular shape. Plaque diameters were calculated, plotted, and analyzed using an electronic counter. Plaque sizes were determined in three independent biological repeats.

### Virus characterization by RFLP-PCR

To confirm the presence of targeted virus and evaluate the changes in the virus genome due to intermittent passage, the harvested fluid was assayed by RFLP-PCR using a specific primer designed for ORF 38 and ORF 54 of varicella-zoster virus (Table 3). The infected cell cultures were removed using a rubber policeman 72 h post-inoculation and pooled in 1.5 mL micro tube. The suspension culture fluid was centrifuged at 8000 rpm for 1 min. Total viral DNA was extracted using Qiagen viral DNA extraction kit (QIAamp DNA Mini Kit, Hilden, Germany) and DNA polymerase Tag (Takara) according to the manufacturer’s instructions. The thermal cycle and appropriate primer sequence are mentioned in Table 3. The amplified fragments were electrophoresed on 2.0% Gel agarose and visualized after staining with safe stain. A panel of two restriction enzymes, PstI and BglI, conducted RFLP. The RFLP reaction mixture was performed in a 32 μL containing 15 μL PCR product, 6 μL buffer, 6 μL deionized water, and 2.5 μl of each BglI (Thermo scientific, Massachusetts, U.S.A.) as well as PstI (Thermo scientific). The mixture was incubated for 16 h in a 37°C water bath. The RFLP product was electrophoresed on a 4% agarose gel. The commercial vaccine (vOKa) was used as a positive control. The size of digestion fragment for various types of varicella is presented in Fig 6.

### SDS-PAGE and Western blotting analysis

Once SDS-PAGE was performed western blotting to detect IE62 proteins of VZV, the proteins were transferred onto a PVDF membrane. The membrane was blocked at room temperature with a blocking reagent (Tris-buffered saline, TBS, containing 5% nonfat milk) for 2 h. It was then incubated for 1 h at 4°C with IE62 antibody (Santa Cruz Biotechnology.Inc; sc-17525) and washed three times with TBS containing 0.05% (v/v) Tween-20. The membrane was exposed to HRP-conjugated anti-goat IgG (Santa Cruz Biotechnology. Inc; sc-2020) second antibody for 1 h, washed four times with gentle shaking, and visualized by DAB (3,3’Diaminobenzidine Tetrahydrochloride: Santa Cruz Biotechnology. Inc; sc-209686) staining [46].

### Indirect immunofluorescence assays for cell infection monitoring

Monolayers of Vero cells (10^**5**^cells per well) in Lab-Tec tissue culture chamber slides (Miles Scientific, Naperville, IL.) were infected with three different passages of VZV local isolate and vOKa separately at a concentration of 30 to 50 particles per well of virus. After 72 h, when CPE became visible, cells were washed twice with PBS. The cells were fixed with methanol 100% for 30 min at room temperature, 100 ul of diluted antiserum (1:100 in PBS) per well was added, with the chamber slides incubated for 4 h at 37°C in a humid atmosphere. Cells were rinsed extensively with PBS and incubated for 1 h at 37°C with 100μl of diluted FITC-conjugated monoclonal antibody (1:700 in PBS). After being washed with PBS, the slides were examined with a fluorescence microscope equipped with a 25X/0.75 PL-Fluotar objective and a filter for light with a wavelength of 450 to 490 nm. Photographs were taken by Motic 2828.

### Quantitative PCR for viral load between different passages

The test was performed by designing a primer set for VZV ORF62 and a probe (FAM) using the ABI Prism 7700. Total cellular DNA was extracted from cell culture harvested fluids using a specific Kit (Genoproof; Cat No ISIN/025 Kit, Czech Republic) containing a protected specific sequence for each Herpes virus such as ORF62 for VZV. The kit generated by an internal control for each virus. The viral gene from infected cells at different passages (4^th^, 9^th^, 17^th^, and 27th) was analyzed via quantitative RT-PCR. The quantity of dye FAM released during amplification was calculated as the values of real time fluorescence. In each reaction, the threshold cycle (Ct) is the value showing the cycle number in which the fluorescence exceeds the threshold [9, 45]. Reactions were repeated three times for each sample, and standard deviations were calculated.

### In vivo testing of local VZV in animal model

The antibody development upon subcutaneous (SC) and intradermal inoculation (ID) was tested regarding high and low passages of local isolated VZV, in 350-450-gram guinea pigs as laboratory animal. There were three guinea pigs in each group. The group were named as: A, B, and D and E injected by four doses, from 30th passages of local isolated viruses. The F group received10^3.0^ CCID_50_ /dose from 5th passages as parent virus. The C^+^ group injected by10 ^3.0^ CCID_50_ /dose of vOka vaccine as a standard positive control. The C^−^ as negative control group received 0.5 mL (1 dose) of the water for injection. The blood samples were taken from animal hearts at four stages. Zero-day was pre-inoculation of the virus, involving the first blood sampling. The other four stages of blood sampling were taken at certain time intervals.

### Sample screening by ELISA

Antibody measurement in the blood serum samples was assayed using standard ELISA test [45, 46]. EUROIMMUN anti-VZV virus ELISA Kit was used to measure antibody titer against varicella virus. The procedure was performed based on manufacturer’s instructions (Euro immune Elisa Kit) for IgG and IgM.

### Skin test

To evaluate the stimulation of cellular immunity due to injection of isolated VZV virus, positive control virus suspension was used for injection of ID with a volume of 0.1 mL, in A, B, D, and E groups. This test was performed through injection in the gluteal region. The injection site was then inspected for 24 hours to a week. The area was marked.

## Acknowledgments

The authors wish to express their appreciations to Dr. Karimi from Mofid Pediatrics Infectious Hospital for his valuable contributions in sampling. The authors also wish to express our appreciations to Dr. Hossein Keyvani for his invaluable technical assistance.

## List of Tables captions

**S1: Table 1.** Summary of serological results of 4 serum samples which the related vesicular lesions used to isolate the virus in this study.

**S2: Table 2.** Primer set and thermal cycle in specific PCR to differentiate VZV from other herpes viruses.

**S3: Table 3.** Viral ORF-62 genes expression levels on MRC-5 and RFSC cell substrate.

## Supporting information

### List of Figures captions

**S1: Fig 1.**
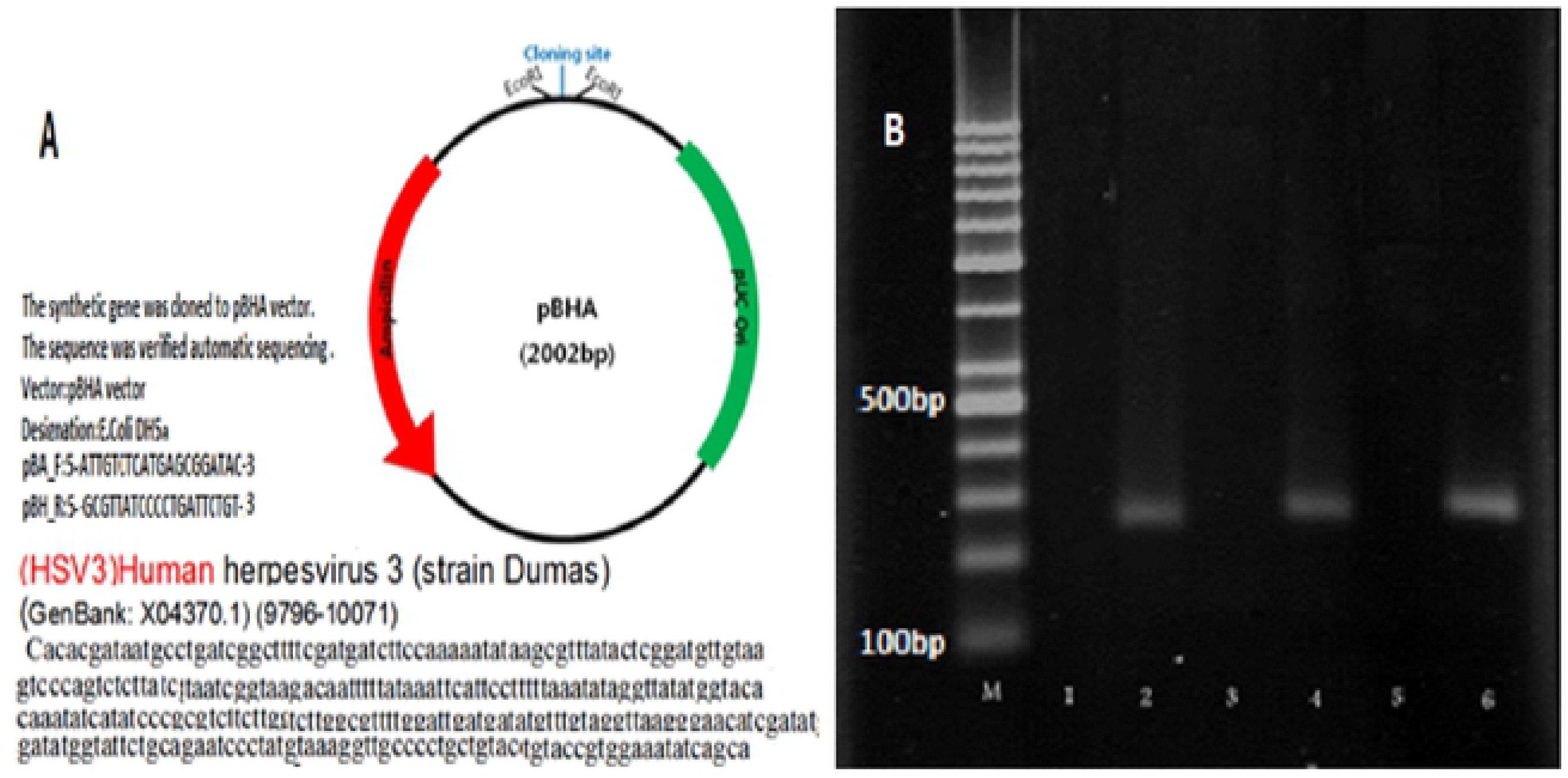
Differentiation of local VZV DNA by specific-PCR. A, pBHA vector; B, Analysis PCR amplification: M lane, Ladder; lane 2, PCR products from control positive (DUMAS STRAIN: 9796-10071) VZV DNA in dilution 1/10; lane 4, PCR products from local VZV DNA in dilution 1/10; lane 6, PCR products from local VZV DNA in dilution 1/100.

**S2: Fig 2.**
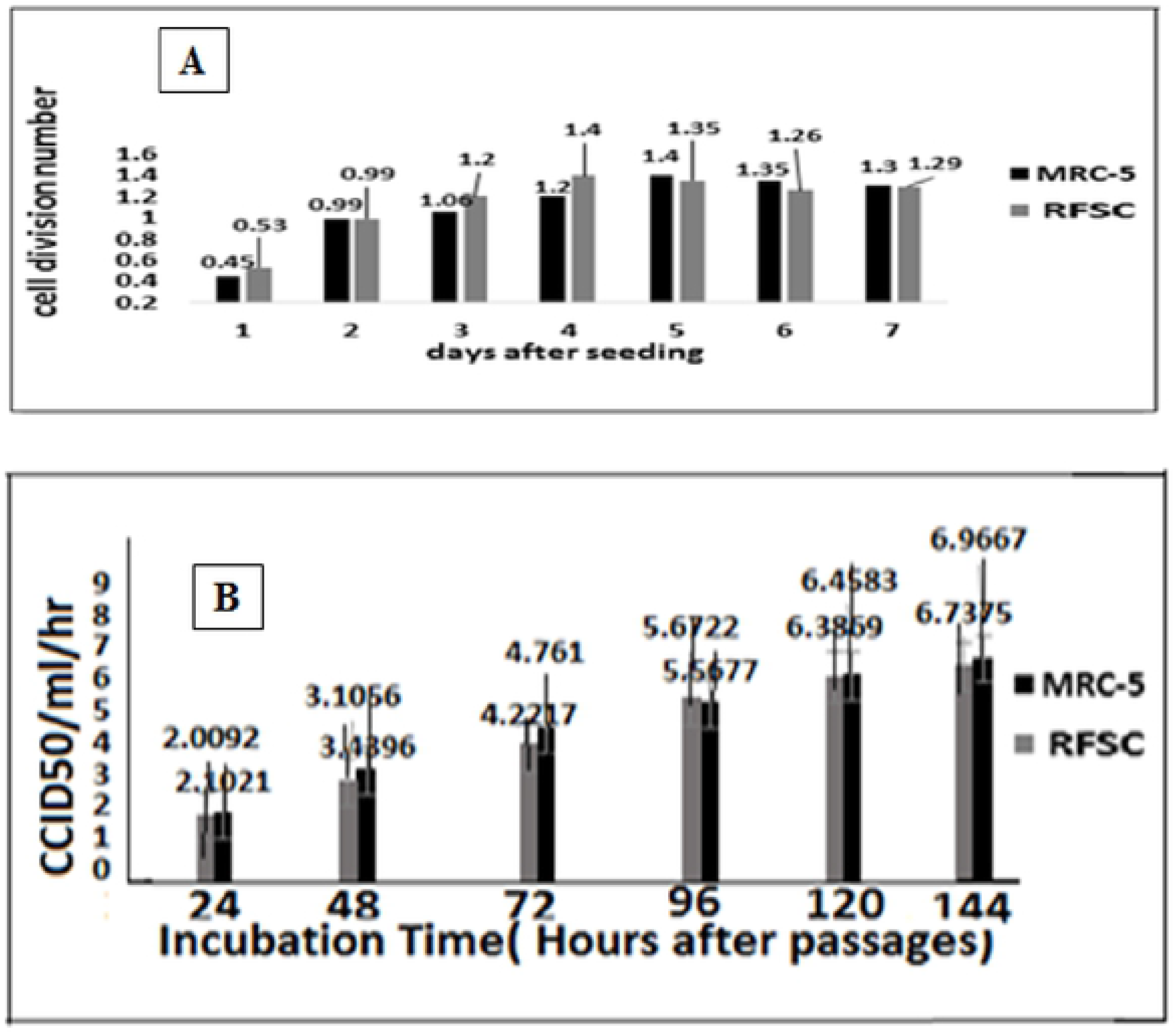
VZV on human diploid lung and human foreskin cells. (A) Comparative cell division number between two human cell substrate during seven days after seeding. MRC-5 as standard human diploid cell strain and RFSC as a continuous local human foreskin cell substrate: The local RFSC at passage 65 and MRC-5 at passages 26. (B) Replication characteristics of local VZV isolate in human diploid lung (MRC-5) and human foreskin (RFSC) cells: Two human cell lines were inoculated with the local VZV and its CPE assessed by CCID_50_ method. Replication kinetics of isolated VZV were compared with RFSC as an alternative cell substrate for future research and production.

**S3: Fig 3.**
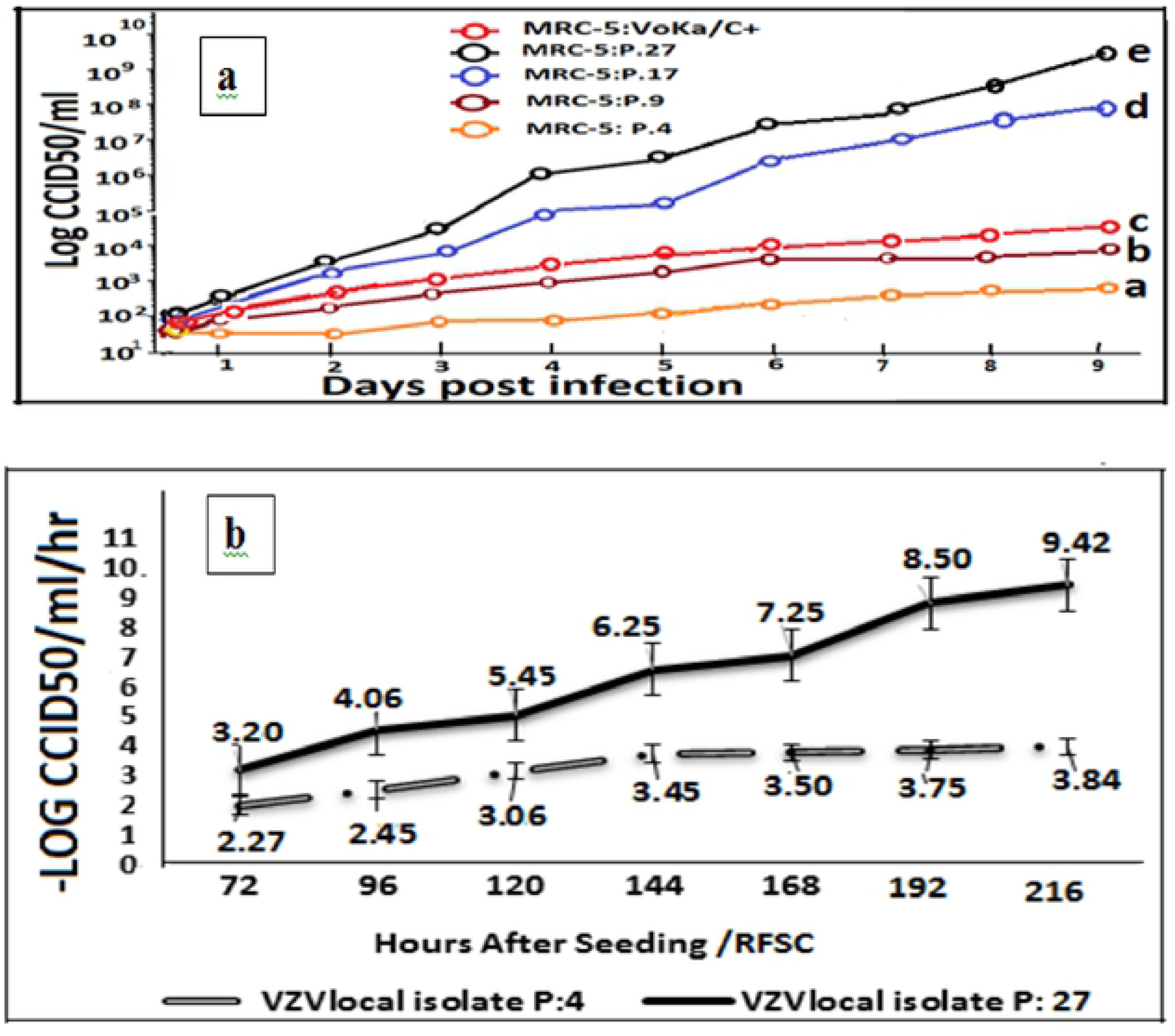
Replication and potency assay of Local isolated VZV virus during different passages on MRC-5. (a) Development of VZV virus inoculated on MRC-5 cell substrate. CCID_50_/mL of VZV virus titers were measured by potency tests as described in the text. The values of, higher and lower level of virus titer was assessed from; p27. (8.75-9.40 LogCCID_50_) and P4. (2.75-3.40 LogCCID_50_). This level for Control positive was (vOKa3.7-4.3LogCCID_50_) after 9 days’ post inoculation. The virus at P.27 and P17 (6.5-7.4 LogCCID_50_) was significant in compare of P9. (3.40-4.5 LogCCID_50_) and P4. (2.75-3.40 LogCCID_50_). Increasing the particles number to 6 and 5LogCCID_50_ for p 27 and P17 was linear to level of virus in RT-PCR but was significant. (b) Replication and potency assay of local isolated VZV during different passages of VZV on local cell substrate (RFSC). After successive serial passages, the virus has become adapted to growing in new cell substrate. This procedure increases the proliferation and number of virus particle. The virus titer has increased by about 6 log CCID_50_/ml in high passages (p: 27), compare to low passages (p: 4) and produces more particle in potency testing at hours post inoculations daily and also after 9 days in high passages The titer of the virus was measured using Karber method.

**S4: Fig 4.**
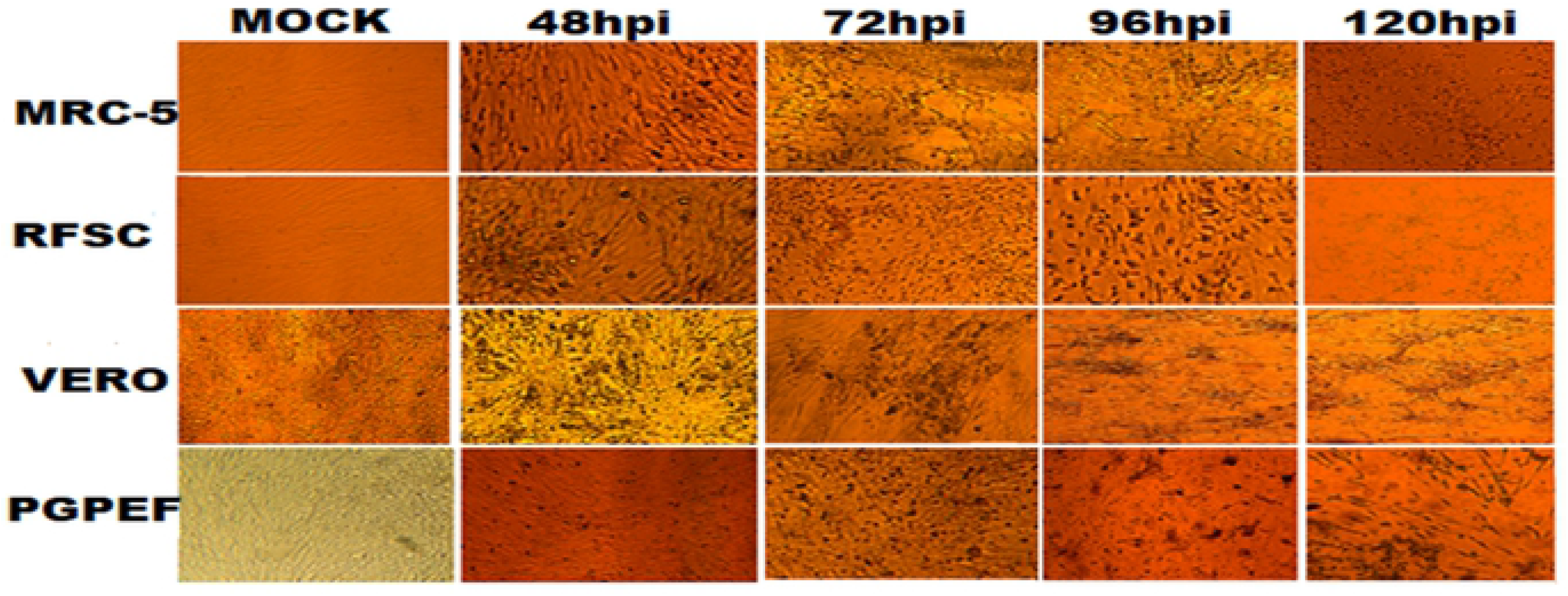
The local VZV cytopathogenicity. Human lung diploid cell (MRC-5) and human foreskin fibroblast (RFSC), Vero, and primary guinea pig fibroblast cells (PGPFC) to local VZV virus infection at 48 and 72and 96 and 120 hours after infection (100x magnification).The virus inoculate as MOI: 0.1 and was in 17th passage.

**S5: Fig 5.**
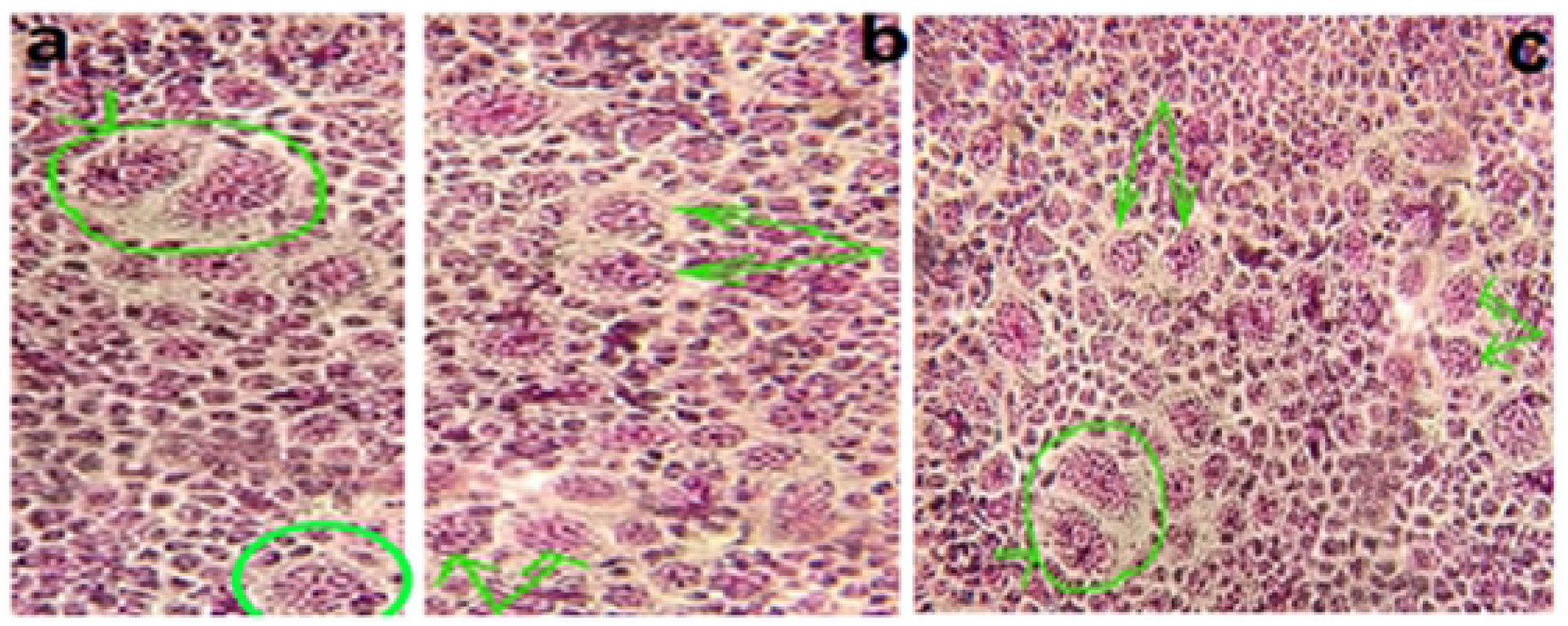
Plaques formed by the viruses that were passaged in cell culture (Scale bar, 200 μ). (a) Representative Images of plaques formed by viruses that were passaged 5 times serially at low multiplicity of infection in cell culture. (b) Representative Images of plaques formed by viruses that were passaged 17 times serially at low multiplicity of infection in cell culture. (c) Representative Images of plaques formed by viruses that were passaged 27 times serially at low multiplicity of infection in cell culture. Serial passaging of viruses has not altered phenotype of plaque of viruses. Although the plaque produced in the 27th passages of virus were larger, and plaques formed by the 4^th^ passages of virus were smaller. But the size distribution of relative plaque diameter normalized against the average plaque diameter of the parental virus. P-values were calculated using multiple comparison test, indicates P<0.05.

**S6: Fig 6.**
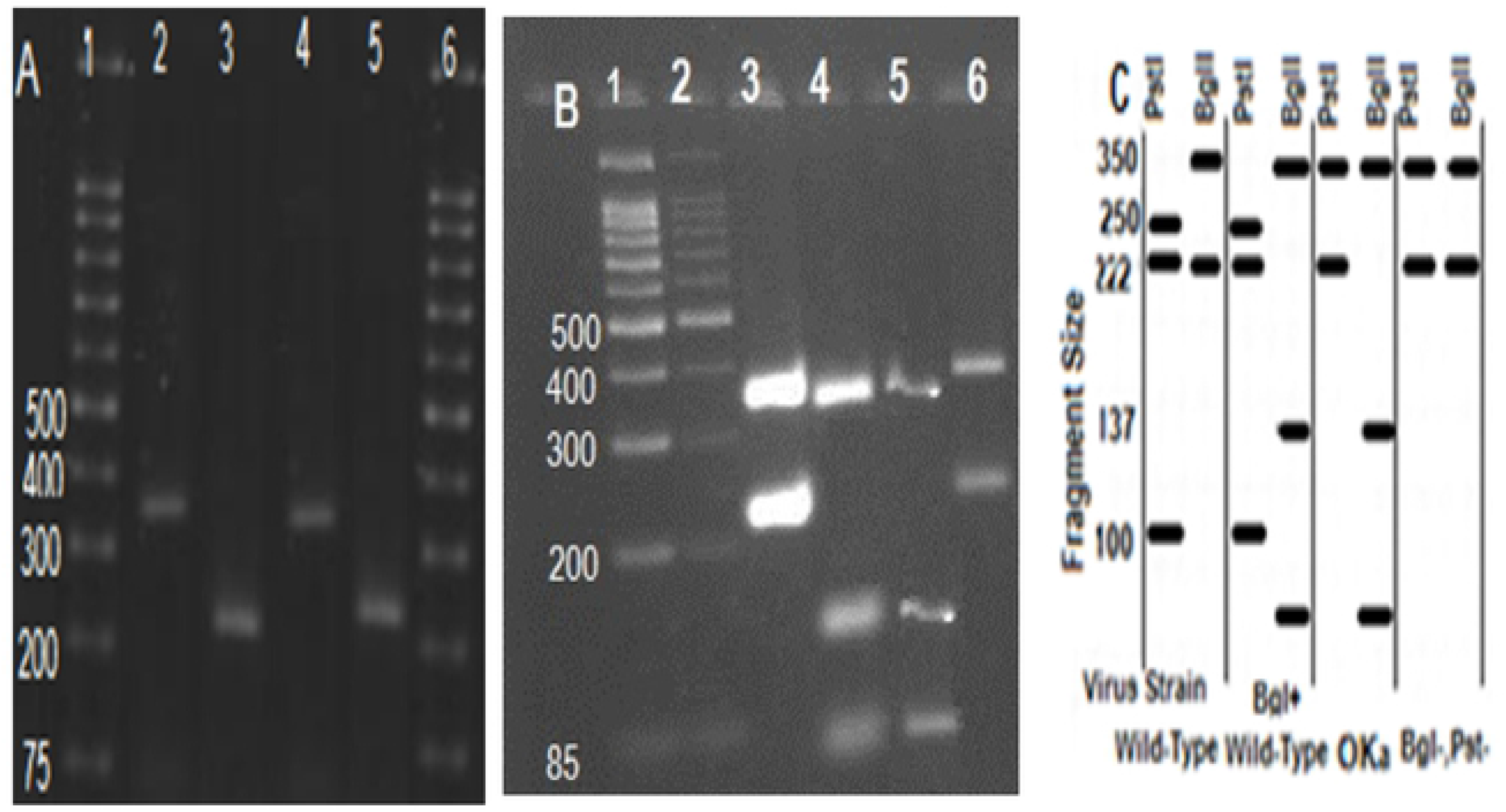
Results of electrophoresis following RFLP test on PCR products of high passage local VZV DNA by RFLP-PCR. (A) PCR amplification of ORF 34 and ORF58, contain 222-bp and 350-bp fragmentlane1, ladder; lane 2and 3, PCR products from vOKa as reference positive, 350-bp and 222-bp; lane 4and5, PCR products from isolated varicella, 350-bp and 222-bp. (B) RFLP on PCR product to analysis the fragment size differentiates wild from attenuated isolated; lane1 and 2 ladder; lane 3, vOKa profile after digestion by PstI (350-bp and 222-bp); lane 4, vOKa profile after digestion by BglI as reference positive (85,100,137bp fragment size); lane 5, digested products from local isolated VZV by BglI (85,100,137bp fragment size); lane 6, digested products from isolated varicella (222 and 350-bp fragment). (C) Schematic RFLP fragment size BglI and PstI.

**Fig 7.**
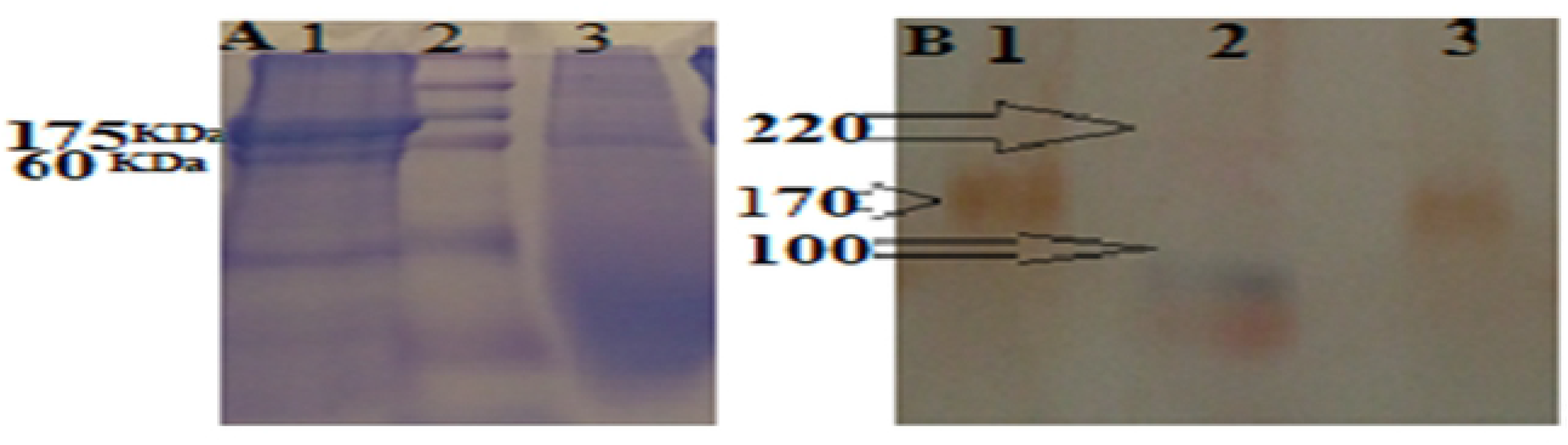
Comparative SDS-PAGE and Western blotting results between isolated VZV and vOKa as control positive. (A) SDS-PAGE of isolated VZV and control positive; Lane 1, 175 kDa band of IE62 protein in the local VZV; Lane 2, Protein Prestain marker; Lane 3, 175 kDa band of IE62 protein in positive control. (B) Western blotting of isolated VZV and control positive; Lane 1, 175 kDa band of IE62 protein confirmed with specific antibody in the local VZV; Lane 2, Protein Prestain marker; Lane 3, 175 kDa band IE62 protein confirmed with specific antibody in positive control. This result confirmed identification of IE62-specific protein as the most important viral transcription activator of VZV gene between isolated VZV and vOKa as control positive. Its presence was examined using labeled specific antibodies, in the Western blot experiment with color band observation. Binding of the labeled secondary antibody was associated with the anti-C-terminal monoclonal antibody near the C-terminal IE62 protein, along with the identification of positive control.

**S8: Fig 8.**
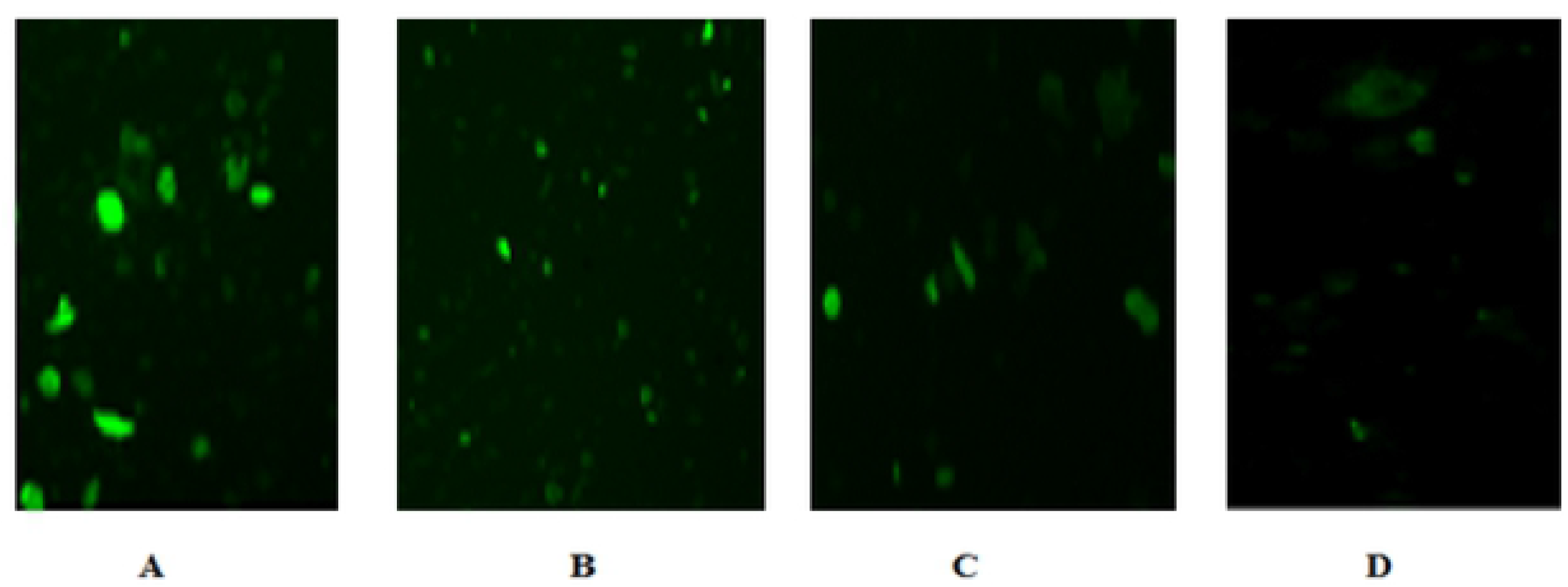
The local VZV in IF test after serial passages. A, the 5th passage of local VZV; B, the 30^th^ passage of local VZV; C, control positive virus; D, the uninfected cell.

**S9: Fig 9.**
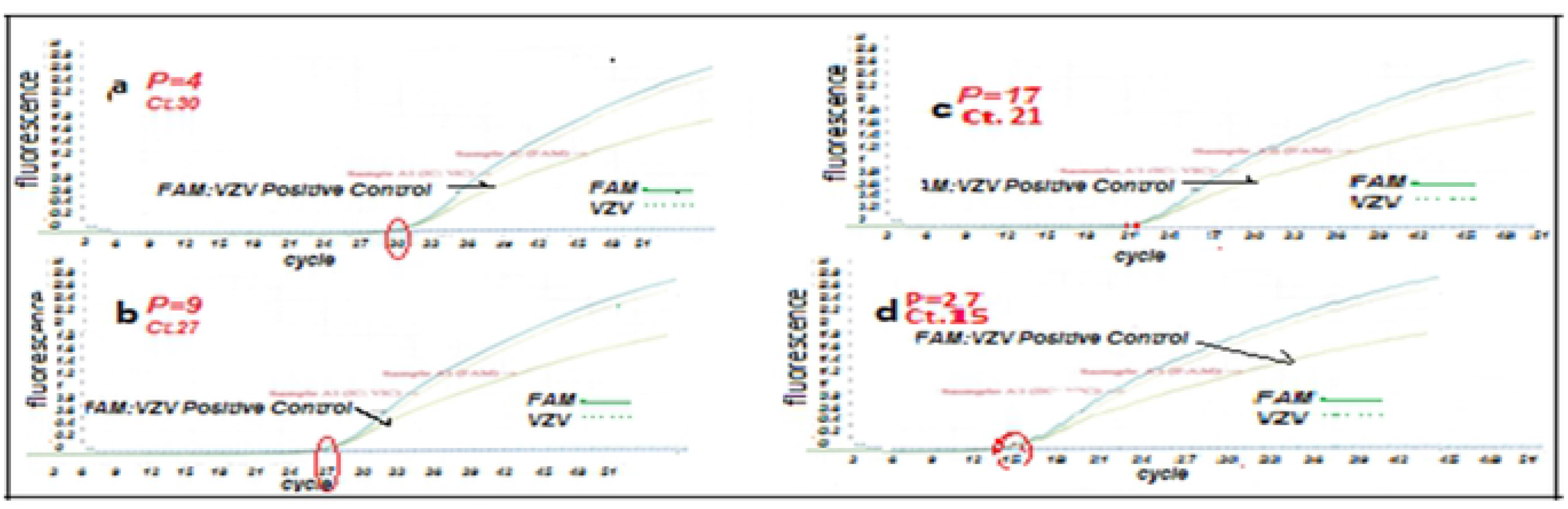
Viral particles in current VZV isolate from Iran by RT-PCR on MRC-5 (72hpi). (a) Isolated VZV-infected MRC-5 cells P.4. (b) Isolated VZV-infected MRC-5 cells P.9. (c) Isolated-infected MRC-5 cells in P.17. (d) Isolated-infected MRC-5 cells P.27.

**S10: Fig 10.**
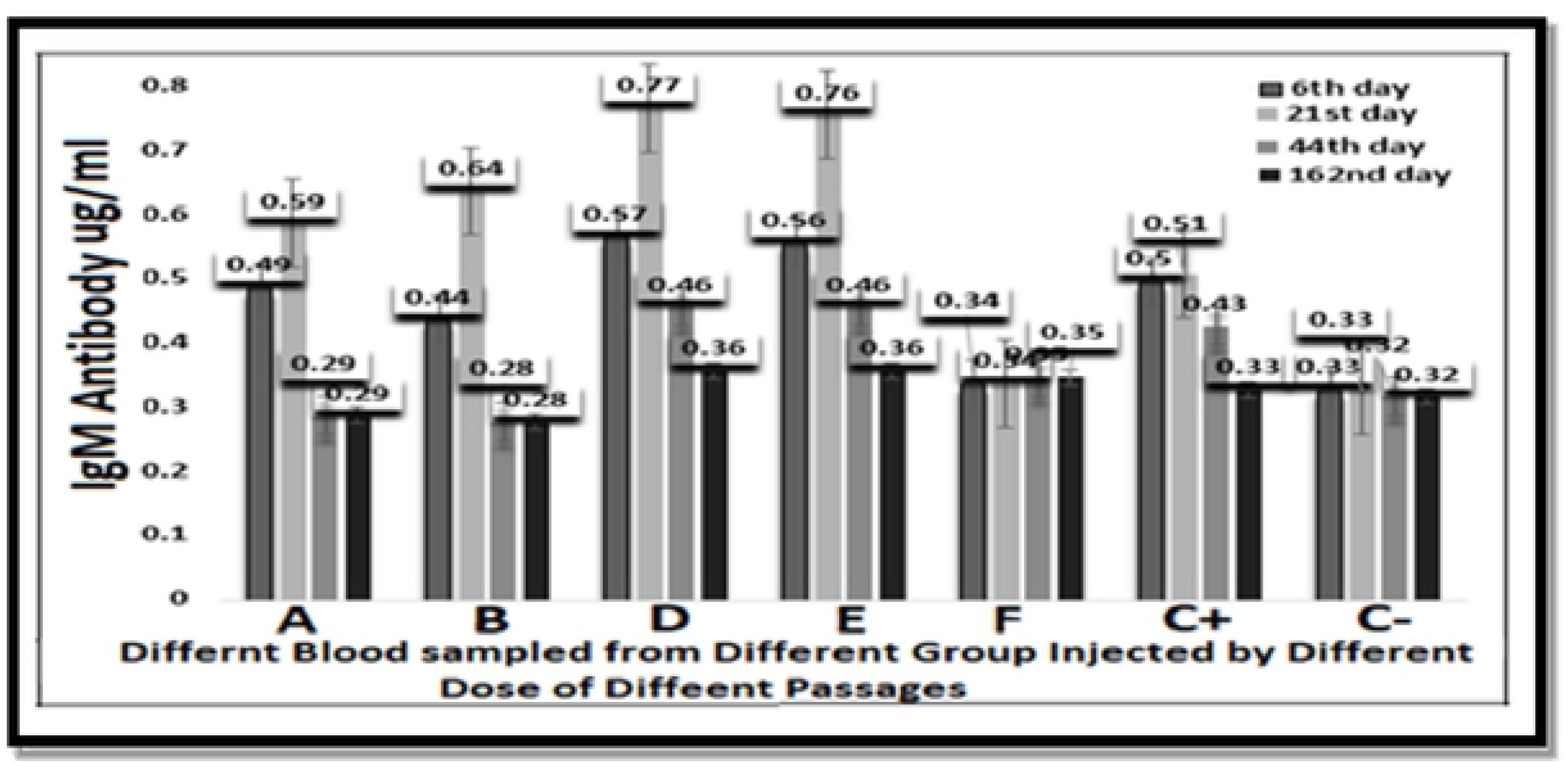
Development of local VZV-specific IgM over time in (350-405 gr) Guinea Pig. Guinea Pig were inoculated ID or SC with (A) 10^5.0^CCID_50_, (B) 104.3 CCID_50_, (D) 10^3.3^ CCID_50_ and (E) 10^3.0^ CCID_50_ of high (30th) passages isolated VZV virus, (F)10^3^ CCID_50_ of low passages (5th) and (C+) 10^3.3^ CCID50 of vOka vaccine as control. The lines indicate threshold level of the assay (P>0.05).

**S11: Fig 11.**
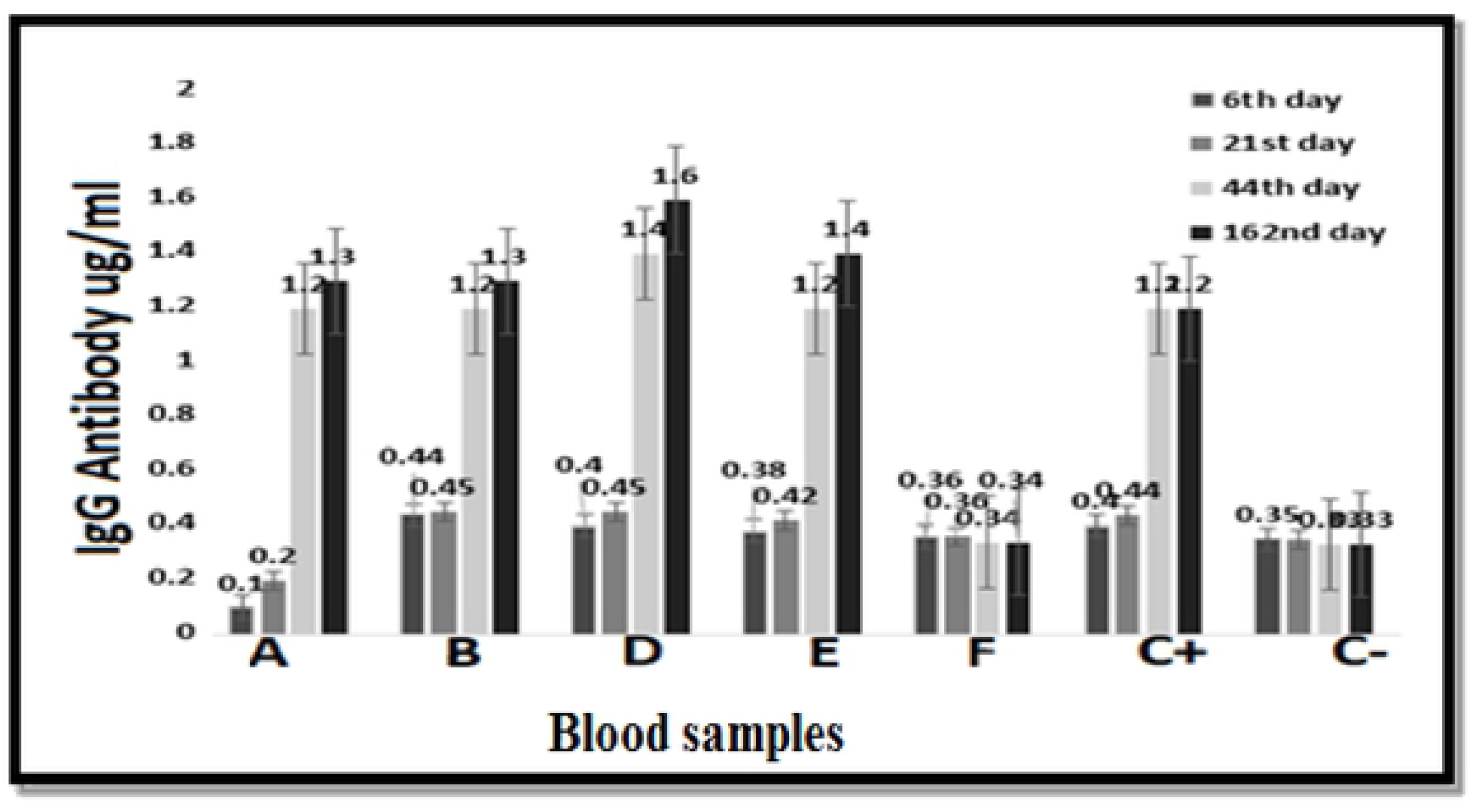
Development of local VZV-specific IgG over time in Guinea Pig (350-405 gr). Guinea Pig were inoculated ID or SC with (A) 10^5.0^CCID_50_, (B) 104.3 CCID_50_, (D) 10^3.3^ CCID_50_ and (E) 10^3.0^ CCID_50_ of 30th) passages isolated VZV virus, (F)10^3^ CCID_50_ of 5th passage and (C+) 10^3.3^ CCID50 of vOKa vaccine as control. The lines indicate threshold level of the assay (P>0.05).

**S12: Fig 12.**
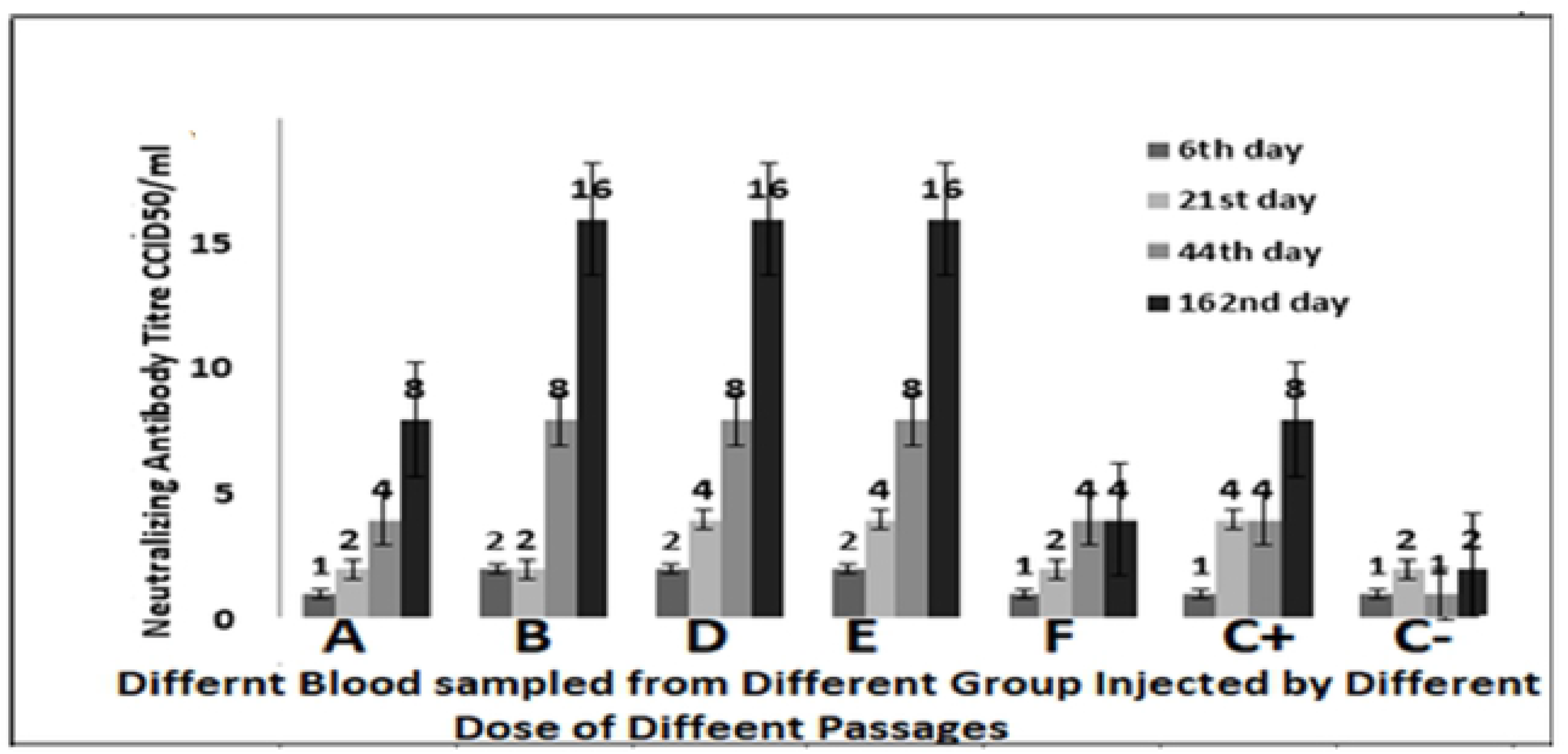
Development of Local VZV-specific neutralizing antibodies over time in (350-405 gr) Guinea Pig. Guinea Pig were inoculated ID or with (A) 10^5.0^CCID_50_, (B) 10^4.3^ CCID_50,_ (D) 10^3.3^ CCID_50_ and (E) 10^3.0^ CCID_50_ of high (30^th^) passages isolated VZV virus, (F)10^3^ CCID_50_ of low passages (5^th^) and (C^+^) 10^3.3^ CCID_50_ of vOKa vaccine as control. The lines indicate threshold level of the assay.

**S13: Fig 13.**
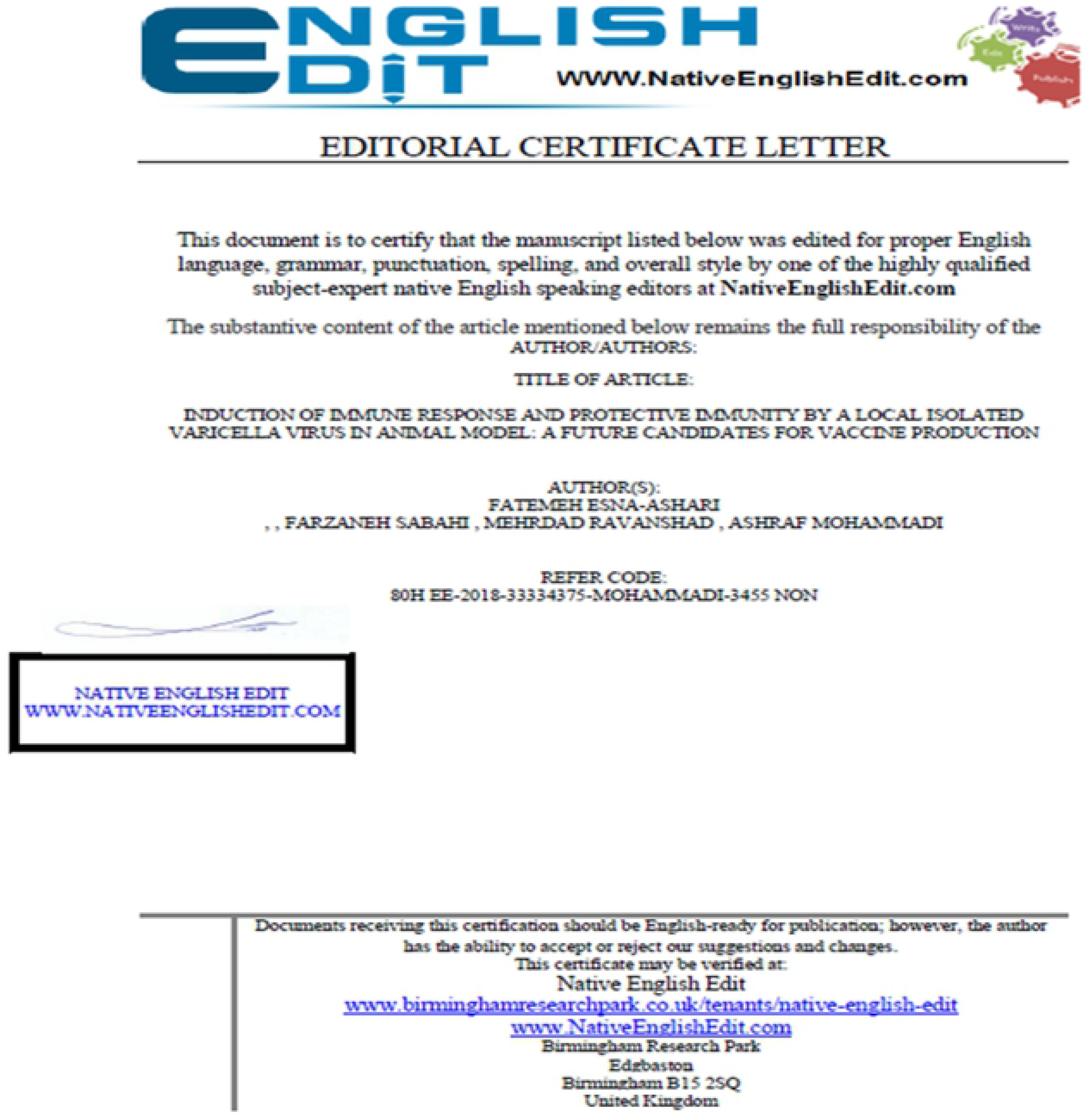
Certification of Native English, Birmingham Research.

## Notes

**Funding:** This work was supported by the Razi Vaccine and Research institute and Tarbiat Modares University as a thesis for Ph.D in medical virology. The funders had no role in study design, data collection and analysis, decision to publish, or preparation of the manuscript.

